# Global landscape of replicative DNA polymerase usage in the human genome

**DOI:** 10.1101/2021.11.14.468503

**Authors:** Eri Koyanagi, Yoko Kakimoto, Fumiya Yoshifuji, Toyoaki Natsume, Atsushi Higashitani, Tomoo Ogi, Antony M. Carr, Masato T. Kanemaki, Yasukazu Daigaku

**Author notes:** These authors contributed equally to this work (Eri Koyanagi, Yoko Kakimoto, Fumiya Yoshifuji). Research Center for Genome & Medical Sciences, Tokyo Metropolitan Institute of Medical Science, Tokyo, Japan (Toyoaki Natsume).

## Abstract

The division of labour among DNA polymerase underlies the accuracy and efficiency of replication. However, the roles of replicative polymerases have not been directly established in human cells. We developed polymerase usage sequence (Pu-seq) in HCT116 cells and mapped Polε and Polα usage genome wide. The polymerase usage profiles show Polε synthesises the leading strand and Polα contributes mainly to lagging strand synthesis. Combining the Polε and Polα profiles, we accurately predict the genome-wide pattern of fork directionality plus zones of replication initiation and termination. We confirm that transcriptional activity contributes to the pattern of initiation and termination and, by separately analysing the effect of transcription on both co-directional and converging forks, demonstrate that coupled DNA synthesis of leading and lagging strands in both co-directional and convergent forks is compromised by transcription. Polymerase uncoupling is particularly evident in the vicinity of large genes, including the two most unstable common fragile sites, FRA3B and FRA3D, thus linking transcription-induced polymerase uncoupling to chromosomal instability.

## Introduction

Accurate DNA replication underlies stable genetic inheritance and is essential in all eukaryotic organisms. In humans, the loss of replication fidelity is responsible for genetic changes that cause both inherited syndromes and somatic diseases, including cancer. There are 16 different DNA polymerases in eukaryotes and the fidelity and efficiency of their synthetic activities are distinct^1^. The division of labour among these polymerases is, therefore, a primary factor in determining the accuracy of genome duplication. In both the budding and fission yeasts, three DNA polymerases: Polδ, Polε and Polα, have been demonstrated to be required for genome replication and are thus termed replicative polymerases. To start all canonical replication events, primase initiates a short RNA primer that is subsequently extended for 10-20 nucleotides by Polα. On the leading strand the bulk of DNA synthesis is subsequently completed by Polε. On the lagging strand, where synthesis is by necessity discontinuous, Polδ takes over from Polα to extend the synthesis up to 100 to 200 bp, generating the Okazaki fragment.

The roles of the replicative polymerases were first established in budding yeast from the mutational bias caused by altered (mutagenic) replicative polymerases in the vicinity of an efficient replication origin, where replication directionality could be predicted^2, 3^. Using a similar mutational bias approach in fission yeast, the role of Polδ in lagging strand synthesis was shown to be conserved. Using a mutated Polε that is prone to introducing ribonucleotides (rNMPs) into DNA it was also shown, using strand-specific alkali sensitivity, that the role of Polε in synthesising the leading strand was similarly conserved ^4^. To expand the analysis of polymerase usage genome-wide, the locations of the increased levels of rNMPs incorporated by individual mutated DNA polymerases (Polα, Polδ or Polε) were identified by whole genome sequencing^5–7^. These data provided direct evidence that the bulk of leading strand synthesis is performed by Polε, while that of the lagging strand was the responsibility of Polα and Polδ. Consistent with these in vivo reports, the roles of budding yeast replicative polymerases have similarly been demonstrated by in vitro studies that reconstituted the replisome with purified factors^8, 9^.

In addition to confirming the division of labour among replicative DNA polymerase, the genome-wide data of replicative polymerase usage also provided highly detailed and discriminatory information about replication fork dynamics. For example, genomic sites with an increased probability of either replication initiation or termination associate with reciprocal changes in leading and lagging strand polymerase usage. By calculating relative changes (differential derivatives) of the profiles of the individual replicative polymerases, the population percentage of replication initiation and termination events were globally measured. This approach identified replication initiation sites, plus their probability of initiation (efficiency), at unpreceded resolution in both budding and fission yeast. It also provided an estimation of the probability of termination across the genome^5, 10^. The accuracy of the methodology was exemplified by the fact that initiation sites identified in budding yeast correspond with the known sequence-specificity of replication origins. In fission yeast, the initiation sites correlated closely with AT richness^11^. Indeed, Monte Carlo simulation of replication fork dynamics based solely on the predicted distribution of the origin recognition complex calculated from the genomic AT content produced a profile of fork dynamics that was strikingly similar to the experimental data^12^.

The profiles of leading and lagging strand DNA polymerase usage can also be directly converted into replication fork directionality (RFD), which represents the proportion of leftward or rightward moving forks at each genomic locus. Again, the accuracy of these data has been verified by multiple studies. For example, mathematical analysis of the RFD data derived from polymerase usage has been used to predict replication timing (RT) across entire chromosomes. The resulting data is superimposable on experimental RT data derived from measured DNA copy number^5^. Given the precision, quantitative accuracy and the concordance with previous measurement of replication fork dynamics, the identification of polymerase usage provides a rational approach to explore genome replication globally in other eukaryotic organisms.

In contrast to lower eukaryotes, the division of labour among replicative polymerases in metazoan cells remains to be addressed. Although human Polα and Polδ have been shown to be required for the synthesis of both leading and lagging strands in reconstituted SV40 replication systems^13^, the usage of replicative polymerases during genomic DNA replication has not been directly characterised. To elucidate the division of labour among replicative polymerases in human cells, and to analyse the profile of replication forks at high resolution, we set out to track the usage of leading and lagging strand DNA polymerases across the human genome. In order to track synthesis by replicative DNA polymerases, we followed the equivalent logic of polymerase usage sequencing (Pu-seq, also known as HydEn-seq) in the yeasts and exploited alleles of replicative DNA polymerases that incorporate an excess of rNMP into DNA during synthesis^5–7^ (**Fig. 1a**).

**Fig. 1.**
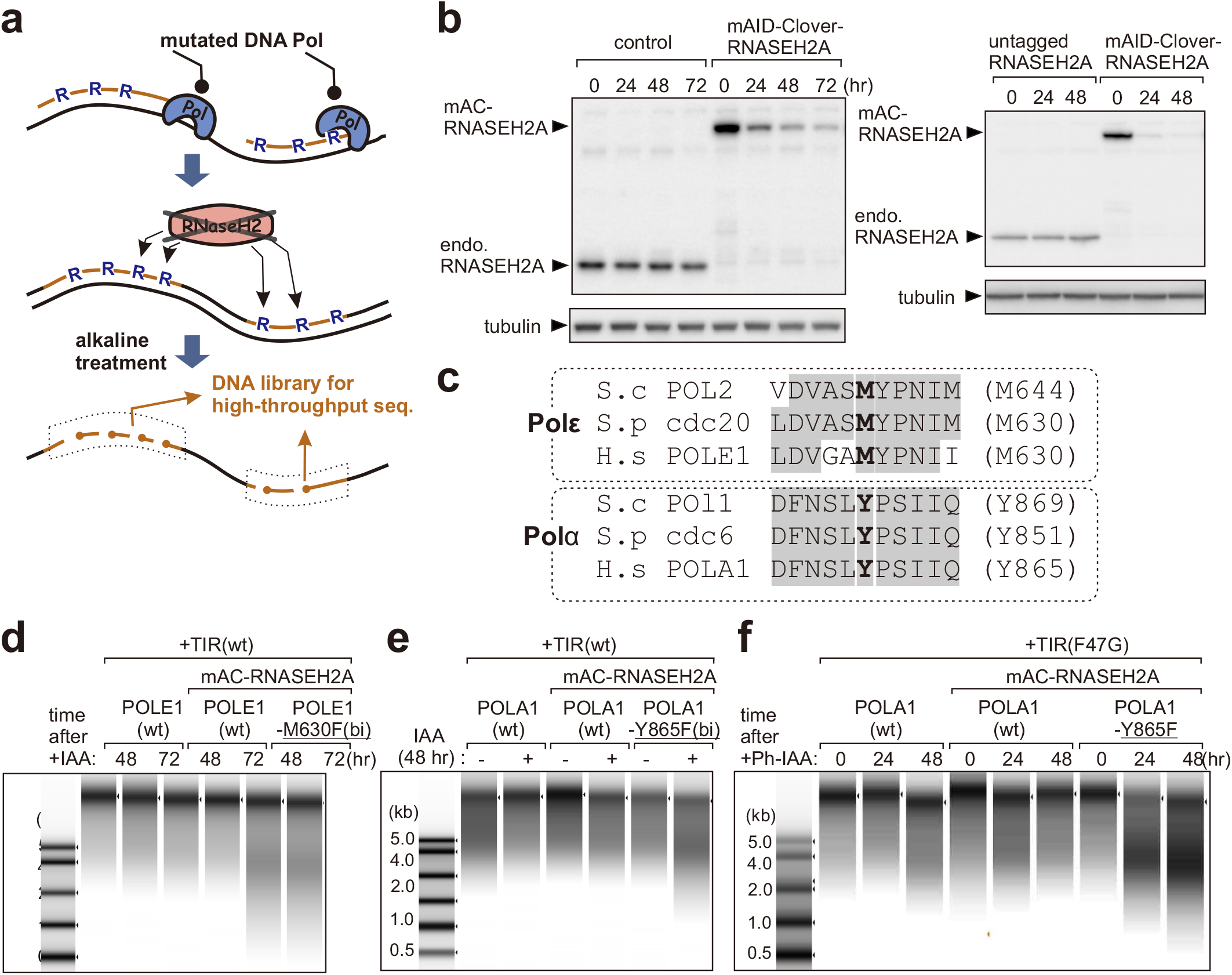
Ribonucleotide incorporation into DNA in POLE1-M630F and POLA1-Y865F cells. **a.** Schematic representation of Pu-seq. Top: ribonucleotides (R) are incorporated by the mutated DNA polymerase. Middle: in the absence of RNase H2-dependent RER, rNMPs remain in the DNA. Bottom: the sugar backbone of DNA strand is cleaved at sites of rNMP incorporation by alkali. Small ssDNA fragments are collected and subjected to library preparation and sequencing. **b**. Left: auxin-induced degradation of RNASEH2A following addition of indole-3-acetic acid (IAA) to cells expressing wild type *O. sativa* TIR1 (AID system). Right: 5-Ph-IAA-induced degradation of RNASH2A in cells expressing *O. sativa* TIR1(F74G) (AID2 system). **c**. Conservation of targeted amino acid residues of DNA Polε and Polα to induce rNMP incorporation. **d**,**e**,**f**. Extracted genomic DNA from the indicated cell lines after the initiation of RNASEH2A degradation treated with alkali to cleave at incorporated rNMPs and analysed by electrophoresis.

Using the near-diploid colon cancer cell line HCT116 we successfully produced genome-wide profiles of Polε and Polα usage that, respectively, reflected leading and lagging strand synthesis. By analysing these profiles we confirm that transcriptional activity influences replication initiation, demonstrate that fork directionality impacts termination close to transcription start sites and show that transcription can perturb the coupling of leading and lagging strand polymerases. Finally, we also show that, at several common fragile sites (CFS) expressed in HCT116 cells, a high level of uncoupled polymerase usage is apparent due to the local inhibition of leading strand DNA synthesis.

## Results

### Construction of ribonucleotide-incorporating DNA polymerase mutant lines

Ribonucleotides are normally incorporated by the replicative polymerases approximately 1:4000 incorporation events. The mutated alleles of replicative polymerases used in yeast to map polymerase usage significantly increase this rate of incorporation. Nonetheless, such rNMPs are rapidly removed from duplex DNA by ribonucleotide excision repair (RER), which is initiated by the RNase H2 enzyme. Therefore, RNase H2 must be inactivated concomitantly with increased rNMP incorporation in order to map the distribution of rNMPs genome-wide. Unlike in yeast, RNase H2 is essential for growth of mammalian cells in the presence of p53^14^. Thus, we developed a HCT116 cell line where we could induce acute degradation of the largest RNase H2 subunit, RNASEH2A, by the addition of auxin. In this auxin-inducible degron (AID) system^15^ the target protein, RNASEH2A, is tagged with the minimal auxin-inducible degron tag (mAID) and an auxin receptor protein from rice (OsTIR) is expressed. Treatment of the cells with the plant hormone 3-indole-acetic acid (IAA) promotes an interaction between OsTIR1 (an Cullin E3 ubiquitin ligase component) and the mAID-tagged RNASEH2A. Recruiting mAID tagged RNASEH2A to the Cullin E3 ligase by the addition of IAA induced ubiquitylation-dependent degradation of the targeted protein (**Fig. 1b** left).

To generate mutant alleles of the POLD1, POLE1 and POLA1 genes (encoding the catalytic subunits of Polδ, Polε and Polα respectively) that are predicted to promote increased rNMP incorporation during synthesis we aligned the highly conserved amino acid sequence of their catalytic sites with the corresponding yeast polymerases. This identified POLD1-L606G, POLE1-M630F and POLA1-Y865F as equivalent mutations to those exploited in budding and fission yeasts. Using CRISPR-Cas9-mediated homology-directed repair, we attempted to generate bi-allelic mutations in the RNASEH2A-degron HCT116 cells (**Fig. 1c**). As a result, POLE1-M630F and POLA1-Y865F mutations were independently introduced into both alleles of the corresponding gene. The POLD1-L606G mutation was, however, only introduced into the one allele, even after repetitive trials. We also observed that ectopically expressing the POLD1-L606G gene does not provide the essential function of the POLD1 gene.

Using the biallelic mutant cell lines for Polε and Polα, in addition to relevant control cell lines, we examined whether RNASEH2A degradation causes increased levels of rNMP in the DNA. Genomic DNA was extracted, treated with alkaline (which preferentially hydrolyses the phosphate-backbone 3’ of the incorporated rNMPs) and the extent of fragmentation was examined by running the denatured samples on agarose gels^5, 6^. For the POLE1-M630F and POLA1-Y865F cell lines, increased levels of small DNA fragments were observed upon RNASEH2A degradation when compared control cell lines. This indicates that rNMPs are incorporated at appreciably higher levels by the mutated Polε and Polα (**Fig. 1de**). However, rNMP incorporation in the POLA1-Y865F cell line was limited in comparison to the POLE1-M630F cell line. We therefore adapted the POLA1-Y865F cell line to the recently developed AID2 system that exploits the highly specific binding of a mutated version of OsTIR (F74G) with an auxin analogue, 5-Ph-IAA^16^. As expected, more efficient degradation of RNASEH2A was observed (**Fig. 1b** right) and considerably increased levels of incorporated rNMP were evident in the POLA1-Y865F cell line when compared to controls (**Fig. 1f**).

### Mapping polymerase usage across the genome

To map usage of Polε and Polα, DNA was prepared from the relevant cell lines 48 hours after IAA/5- Ph-IAA addition and small alkaline-cleaved single stranded-DNA (ssDNA) fragments (< 2kb) were collected and used to produce libraries for Illumina sequencing. Approximately 200 million paired-end reads were obtained for each cell line and the positions of 5’ ends were mapped to either the Watson or Crick strands. The 5’ ends represent the rNMP positions, which were scored in 1-kb bins across the genome. The relative ratio of reads for Polε and Polα mutants, when compared to those of control lines, provides scores representative of relative usage of these polymerases (**Fig. 2a**). Along the chromosome coordinates, a reciprocal relationship was evident between the profiles of Polε and Polα on the same strand. Similarly, a reciprocal relationship was evident between the profiles of same polymerase when comparing the Watson with the Crick strands. These patterns of polymerase profiles are evident across the genome, consistent with the primary roles of Polε in leading strand and Polα in lagging strand synthesis (**Fig. 2b**). Importantly, two independent experiments were confirmed to yield nearly identical polymerase profiles (**Supplementary fig. 1**). These results demonstrate the roles of Polε and Polα are conserved between yeasts and humans, although replicon size (the region replicated from a single replication initiation site) is quite different: 30-50 kb in yeasts vs. several hundred kb to 1-1.5 Mb in humans. Interestingly, visual inspection of the profiles shows that the typical enrichment of either leading or lagging strand synthesis was not evident in some areas of the genome. These regions exclusively locate at heterochromatic late replicating segments (71 sites across the genome, **Supplementary fig. 2abe**). This suggests that DNA replication is regulated differently in these regions (see below).

**Fig. 2.**
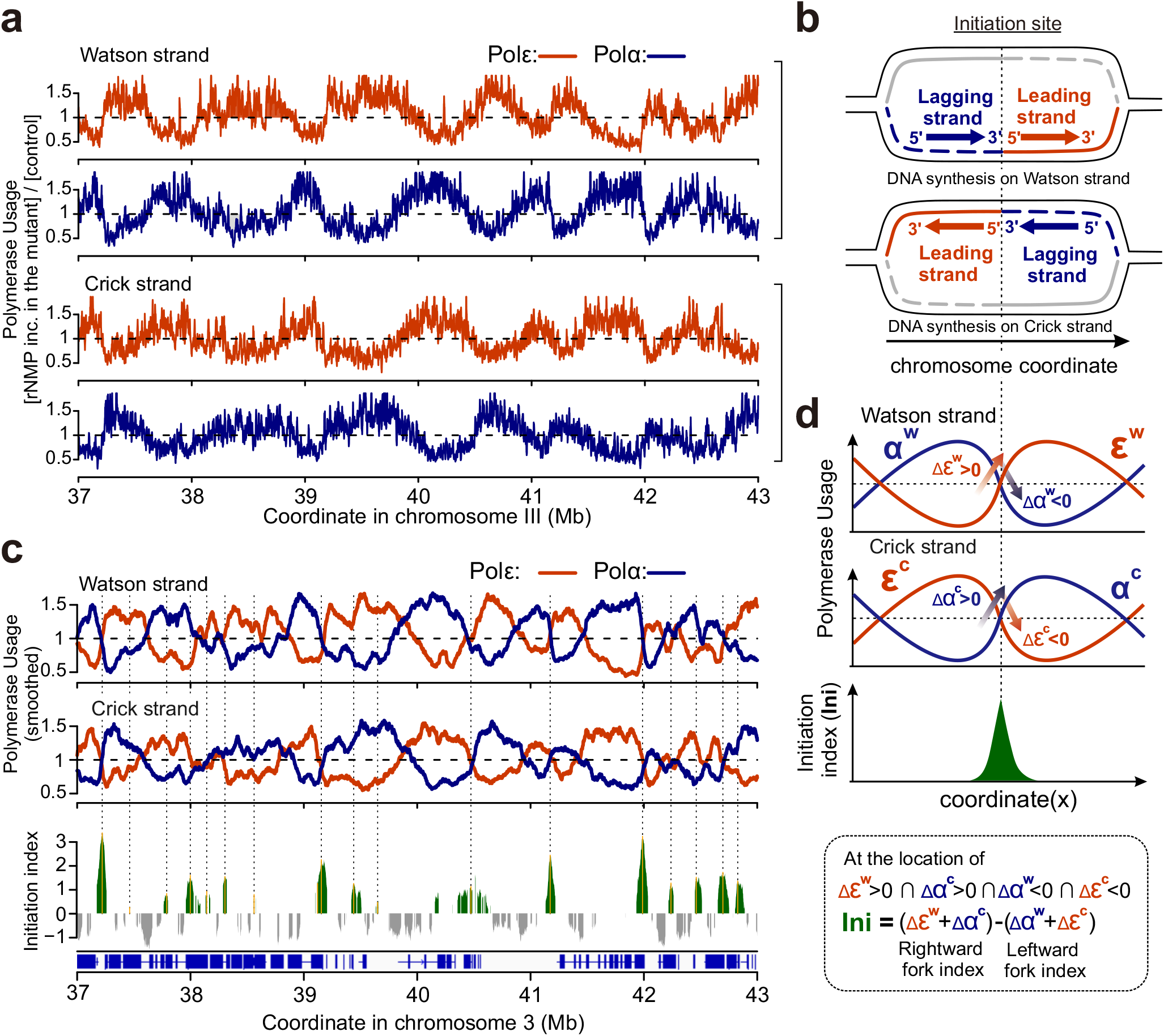
Polymerase usage and replication initiation across the human genome. **a**. Profiles of the relative reads for Polε and Polα mutants on the Watson and Crick strand for each 1 kb bin. Orange: Polε (POLE1-M630F). Blue: Polα (POLA1-Y865F). A representative region of chromosome 3 is shown. Data were smoothed with a moving average (m = 3; see materials and methods). **b**. Schematic representation of predicted Polε and Polα profiles at a site of replication initiation. Orange: leading strand. Blue: lagging strand. **c**. Top: smoothed data from panel a (moving average, m = 30) provides a map of polymerase usage. Bottom: plot of the calculated initiation index. Positive values (green) represent increased initiation activity. Negative values (grey) represent increased converging fork termination. **d**. Definition of the initiation index (see materials and methods for further details). ’Δ’ indicates the differential between neighbouring bins, e.g. at location x, Δε(x) is defined as ε(x+1)-ε(x).

The profiles of leading and lagging strand polymerases provide two direct and independent measurements of the proportions of replication forks moving either rightward or leftward at each location across the genome. We therefore calculated replication fork direction profiles independently from either the Polε (RFD^ε)^ or Polα (RFD^α^) data and compared these with OK-seq replication directionality data that we calculated from sequencing data^17, 18^ using the identical algorithm (RFD^OK^; for details see materials and methods). Visual inspection indicates that the overall trends of RFD^ε^, RFD^α^ and RFD^OK^ are highly similar (**Supplementary fig. 3ab**), albeit with the Pu-seq derived data showing lower amplitude peaks. RFD^α^ is more similar to RFD^OK^ than to RFD^ε^ (**Supplementary fig. 3c**), indicating that the profile of Polα captures signatures specific to lagging strand synthesis. Considering the minor differences between leading and lagging strand synthesis, we established a combined RFD profile from pooled Polε and Polα Pu-seq data. Fluctuation outside of the local trends are notably reduced when compared to RFD^ε^ or RFD^α^ (**Supplementary fig. 3a)**. Thus, combining both leading and lagging strand profiles improves precision as well as resolution of the replication fork profiles (RFD^ε|α^).

**Fig. 3.**
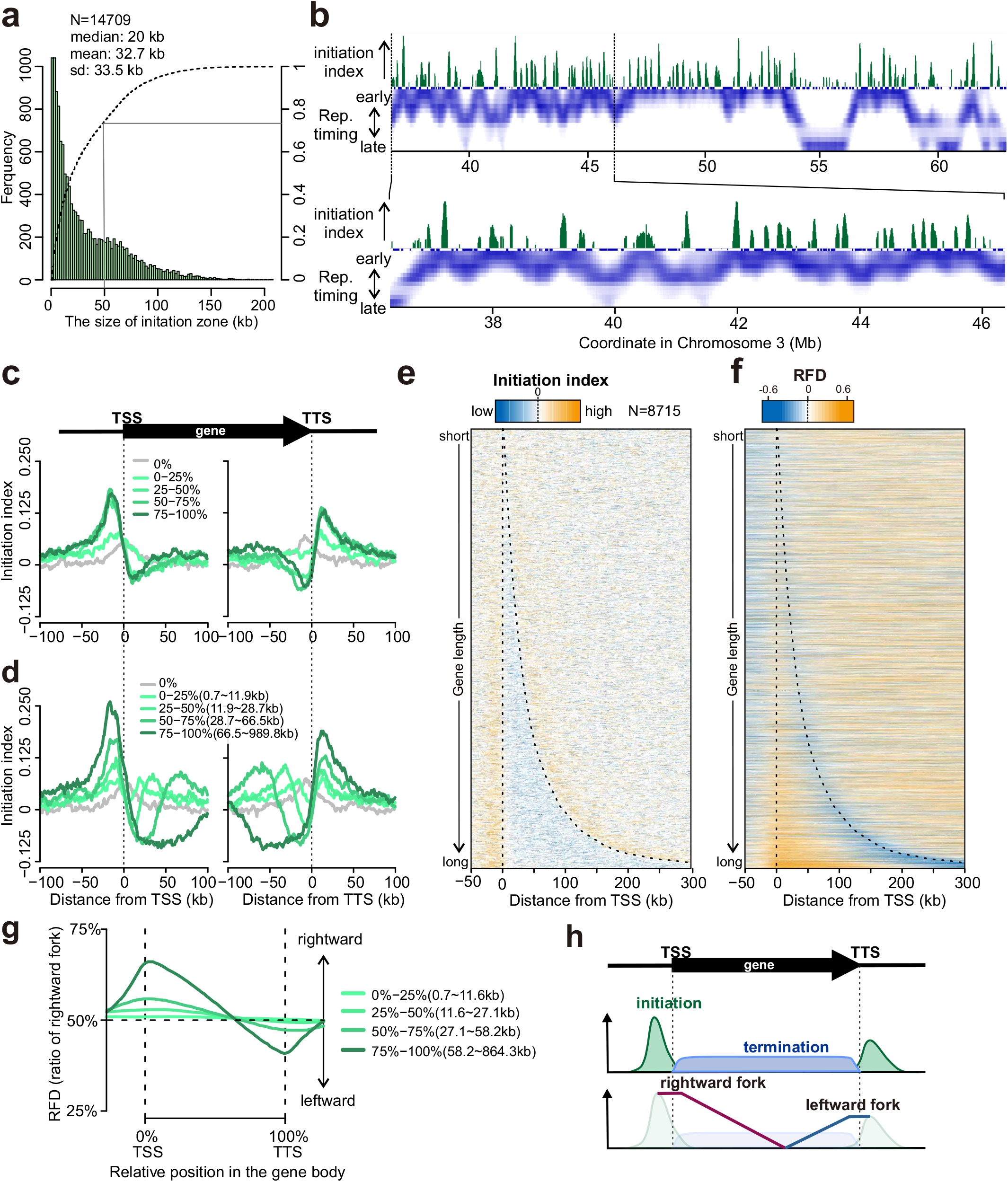
Genomic distribution of initiation sites. **a**. Distribution of initiation zone widths. **b**. High-resolution replication timing data and initiation index plotted for a representative region of chromosome 3. The profile of replication timing for HCT116 cells is from Zao et al (2020)^21^. **c**,**d**. Average initiation index +/- 100kb around annotated TSS and TTS in the human genome. Initiation index data are categorised by transcriptional activity (c) or gene length (d). For the gene length analysis only the 50% most transcriptionally active genes were included. **e**. Heat map representation of data in panel d sorted by gene length. Broken lines indicate the position of TSS and TTS. **f**. Equivalent heat map representation of RFD aligned at TSS and TTS. **g**. Average RFD for the relative positions from TSS to TTS for all genes scaled to the same arbitrary length. RFD values from Supplementary Figure 2 were converted to rightward fork proportion. **h**. Schematic representation of initiation and termination of replication forks as well as fork directionality around a representative gene.

### Defining replication initiation regions from polymerase usage

The division of polymerase labour between leading and lagging strands dictates that sites of frequent replication initiation manifest as reciprocal demarcations in Polε and Polα usage. Therefore, we defined an initiation parameter to represent the local activity of initiation events (**Fig. 2c**). Specifically, we calculate two independent initiation indices (**Supplementary fig. 4ab**), one from the Polε data (Ini^ε^, where Polε synthesis increases towards the 3’ on the Watson strand and Polε decreases towards the 5’ on the Crick strand) and a second from the Polα data (Ini^α^, where Polα synthesis decreases towards the 3’ on the Watson strand and Polα increases towards the 5’ on the Crick strand). These two independent indices can be plotted separately (**Supplementary fig. 4ab**) or combined into a more accurate and constrained cumulative initiation index (Ini^ε|α^, **Fig. 2cd**). The two replicates of the Pu-seq experiments were used to generate two independent combined initiation indices (**Supplementary fig. 4c**). Positive peaks in the initiation index are interpreted as replication initiation sites and peak height as proportional to the population frequency of initiation. Approximately 12,000 initiation peaks were detected in each replicate. The concordance in the peak positions between the two replicates increases with increasing values of the initiation index; when all the peaks were taken into account, 67.2 % of all peaks colocalise, whereas this number increased to 87.5% with top 50% of initiation peaks. (**Supplementary fig. 4d**). Since polymerase profiles were derived from asynchronous cells, this analysis detects initiation events during the entire S phase. Of note, the initiation index can also have negative values, which represents regions where termination of merging forks is frequent.

**Fig. 4.**
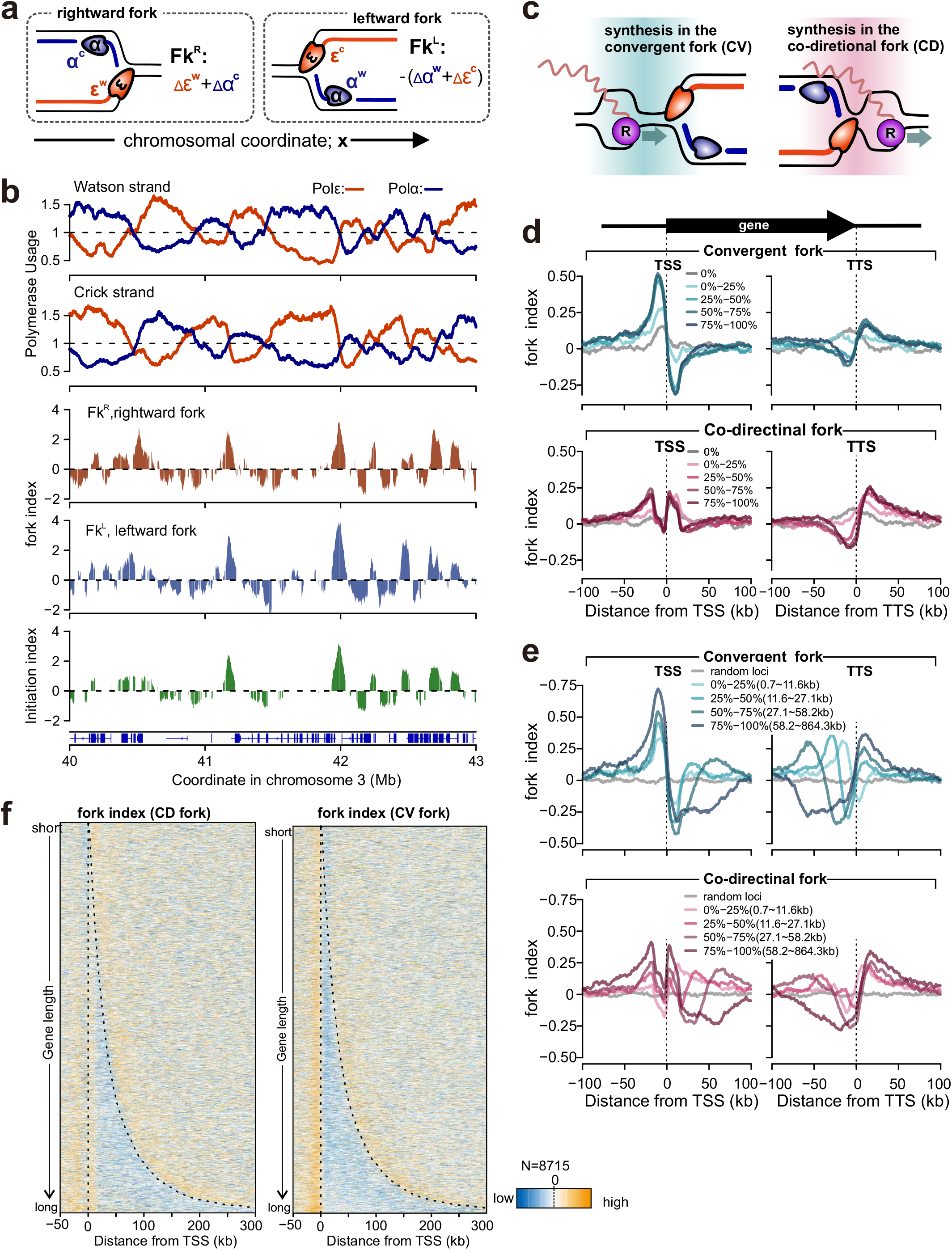
The profiles of rightward and leftward forks. **a**. Definition of fork indices of rightward leftward forks (Fk^R^ and Fk^L^, for further details see materials and methods). ’Δ’ indicates the differential between neighbouring bins, e.g. at location x, Δε(x) is defined as ε(x+1)-ε(x). **b**. Profiles of polymerase usage (top), fork index (middle) and initiation index (bottom) for a representative region of chromosome 3. **c**. Schematic representation of transcription and replication conflicts for convergent (CV) or co-directional (CD) forks. **d**,**e**. Averaged fork index +/- 100 kb around annotated TTS in the human genome. Data for the fork indices of CV and CD forks are categorised by transcriptional activities (d) or gene length (e). For the gene length analysis only the 50% most transcriptionally active genes were included. **f**. Heat map representation of data in panel e sorted by gene length. Broken lines indicate the position of TSS and TTS.

Previously mapped initiation sites in the human genome showed that a proportion localised to a few kilobases, whereas other mapped to ‘initiation zones’ of ∼10-50 kb^17, 19, 20^. In our initiation index profile, the width of positive peaks averaged ∼33 kb (replicate 1: 32.7kb, replicate 2: 33.1 kb) with more than 20% above 50kb (**Fig. 3a**) confirming that initiation events in human cells cluster in zonal regions. The activity of initiation per zone size increases with size, up to approx. 70 kb (**Supplementary fig. 4e**) and, when the initiation index is plotted with high-resolution RT^21^, the zones consistently locate at local peaks of RT (**Fig. 3b, Supplementary fig. S5**). We note that initiation zones are present both in mid-late RT regions (e.g. around 41Mb, 60 Mb in Chr. 3 in **Fig. 3b**, locations marked by circles in **Supplementary fig. S5**) as well as early replicating regions. These data thus demonstrate that high probability initiation zones are not located primarily in early replicating regions, but exist in mid-late replication regions. This is contrary to the prevailing view that efficient initiation is the predominant determinant of early replicating regions and indicates that late-firing but efficient initiation zones exist across the human genome.

We also observed 71 regions of Mb length heterochromatic late replicating regions where clear peaks for initiation zones were not evident (**Supplementary fig. 2c**). These defined the same regions noted above as having unusual fork direction profiles (**Supplementary fig. S2a**). The lack of defined initiation zones was consistent with the reported profile of high-resolution RT^21^, where defined initiation sites were not also observed within these regions (**Supplementary fig. 2d**). This likely reflects that replication initiation occurs at random locations within late replicating heterochromatinic regions. This would also account for the equal frequency of leftward and rightward moving forks.

### Association of replication Initiation/termination with transcriptional activity

A positional relationship between replication initiation and transcription has been highlighted by multiple studies using different techniques^17, 18, 20, 22^. We therefore analysed how the distribution of transcription units influences the initiation and termination of replication forks in our Pu-seq derived data. The initiation index was aligned at transcription start sites (TSS) and transcription termination sites (TTS). The initiation index score increases in the vicinity of both TSS and TTS **(Fig. 3c-e)**. High levels of gene expression (**Fig. 3c**) and increased gene length both correlated with increasing initiation index score (**Fig. 3d**), consistent with published OK-seq data from RPE-1 cells^18^. However, the higher resolution of the Pu-seq derived data shows that the peak of initiation localises ∼20kb upstream of the TSS and ∼20kb downstream of the TTS. We also note that initiation index shows a negative value throughout gene bodies, consistent with frequent termination in these regions (**Fig. 3de**).

We next aligned the Pu-seq-derived RFD data with TSS and TTS. At TSS and the 5’ regions of genes we observe a significant bias of rightward moving forks. At the TTS and 3’ regions of genes we observe a bias towards leftward moving forks (**Fig. 3fg**). Thus, as expected when initiation is biased towards TSS and TTS (and is largely absent from gene bodies), termination is increased within the gene body (**Fig. 3h**). Notably, these localised initiation and termination patterns are far more apparent for large genes. While the distribution of genes is a factor in the location of initiation and termination zones our result also demonstrate that there are many genes that are transcriptionally active but do not show replication initiation in the vicinity of their TSS and TTS. Plotting the extent of replication initiation at TSS/TTS against transcriptional activity, it is evident that most of the genes associated with replication initiation are transcriptionally active. However, transcriptionally active genes are not necessarily associated with replication initiation: approximately 20% of genes with > average transcriptional activity show no evidence of initiation in the upstream of TSS (**Supplementary fig. 6a**). These data suggest that many transcriptionally active genes are passively replicated and transcriptional units do not account for all replication initiation. Furthermore, approximately 60% of initiation zones do not overlap with upstream regions of TSS or downstream regions of TTS (**Supplementary fig. 6b**).

It is evident that more forks travel from the initiation zones upstream of the TSS into the gene bodies (i.e. co-directional with transcription) than travel from the gene body through the TSS convergent to the direction of transcription. However, this is not absolute and we estimate for the top 50% of transcribed genes this equates to between 66-51% co-directional fork and between 49-34 % of forks that are convergent with transcription, which varies dependent on gene length (**Fig. 3g**). To separately visualise the dynamics of replication forks dependent on their orientation at any one locus, we calculated a separate ‘fork index’ for both rightward (Fk^R^) and leftward (Fk^L^) moving forks that independently represent their cumulative initiation and termination behaviours. These profiles can be interpreted as separate ‘initiation indices’ for rightward and leftward moving forks. Thus, fork initiation and termination are represented by positive or negative values (**Fig. 4a**). Visual inspection of genome-wide Fk^R^ and Fk^L^ profiles showed similar genomic profiles that are, as expected, congruent with the initiation index discussed above (**Fig. 4b** the experimental replicate in **Supplementary fig. 7**).

Aligning Fk^R^ and Fk^L^ at TSS allowed us to separately visualise the effect of transcription on forks that are either co-directional (CD) or convergent (CV) with transcription (**Fig. 4cde**). In these figures, leftward moving forks that initiate upstream of TSS – high peak of fork index – move away from the gene and the fork index thus declines slowly. However, leftward moving forks within the gene body are CV with transcription and showed dramatically decreased fork index downstream of TSS (**Fig. 4def**). This suggests that head-to-head transcription replication clashes slows fork processivity and increases termination events in this region. In the case of CD forks (rightward moving) the fork index profile at TSS is more complex: two lower peaks are evident in the vicinity of TSS. We interpret this as a combination of initiation and termination: i.e. a negative signal (fork termination) at and immediately upstream of TSS is embedded within strong positive signals derived from initiation events associated with the upstream 20 kb. It should be noted as discussed above that not all transcriptionally active genes are associated with an increase in initiation upstream of TSS (**Supplementary fig. 6a**). In the transcriptionally active genes (> average) which do not show an associated increase of replication initiation at TSS, the fork index for CD forks drops sharply below zero at and immediately upstream of TSS (‘a-1’ in **Supplementary fig. 6c**), indicating that forks terminate in this region. As expected, the profile of genes that show association of initiation with the TSS resembles that of all active genes (‘a- 2’ in **Supplementary fig. 6c, Fig. 4d**). Thus, this mixed signal of fork termination and initiation manifested in fork index does not necessary represent the profile of individual loci. The combined pattern of CV and CD forks is consistent with the trend for fork initiation in both orientations ∼20 kb upstream of TSS, with the processivity of leftward moving (CV) forks being reduced immediately downstream of TSS (**Fig. 4d** top) and the processivity of rightward moving (CD) being reduced immediate upstream of TSS (**Fig. 4d** bottom). Taking gene length into account demonstrates that this effect is largely independent of gene length and manifests throughout the length of the transcription unit (**Fig. 4e,f**). This tendency is contrary to fork initiation, which increase with gene length, indicating that fork termination also commonly occurs in the vicinity of TSS. During transcription initiation RNA Polymerase II (RNAPII) promoter-proximal pausing is enriched in the immediate vicinity of TSS^23^. Thus RNAPII, directly or indirectly, likely causes an impediment to fork progression. In contrast, as predicted by the initiation index around TTS, both initiation and termination events are evident and fork orientation (co-directional or convergent) did not influence the profiles.

### Local genomic features at replication initiation sites

To investigate if specific sequence-based or chromatin-based features correlate with replication initiation sites, peak positions within initiation zones were computationally identified (yellow vertical lines in **Fig. 2c** bottom and **Supplementary fig. 2c**) and those present in initiation zones of both experimental replicates selected. Using this dataset we examined if specific genomic elements are enriched at these loci. GC skew, AT skew and CpG islands were not enriched. Potential guanine quadruplex (G4) structures were modestly enriched at peaks of initiation index (**Supplementary fig. 8a**). For chromatin features the H2AZ histone variant and, to a lesser extent, trimethylated H3K27 (H3K27me3) were enriched at initiation index peaks (**Supplementary fig. 8b**). In contrast, trimethylated H3K36 (H3K36me3) tended to be excluded from initiation zones and moderately enhanced in flanking regions, likely because it is associated with gene bodies^23^. H3K4me3 was enriched ∼ 20kb either side of the peak of initiation, consistent with its enrichment at TSS^23^(**Supplementary fig. 8c**). Furthermore, analysing genomic status bin based on multiple chromatin profiles by using ChromHMM algorism^24^, initiation zones and TSS/TTS were shown to associate with distinct chromatin status (**Supplementary fig. 8d**). Thus, the chromatin features H2AZ and H3K27me3 positively correlate specifically with initiation sites, while H3K36me3 shows a minor anti-correlation.

To establish which of the features discussed above correlate best with initiation activity we partitioned each chromosome into 10 kb-bins and used principal component analysis (PCA) to deconvolve genomic features in relation to replication initiation (**Supplementary fig. 8e**). This revealed that chromatin-based features (H2AZ and H3K27me3) correlated more closely with replication initiation than G4. As expected, H3K36me3, which is deposited in transcribed regions in an RNAPII-dependent manner (thus marking gene bodies), positioned oppositely to initiation. These data suggest that, combinatorially with transcription, chromatin status is crucial for shaping replication initiation.

### Uncoupling of leading and lagging strand polymerases

Having two independent datasets that represent leading (Polε) and lagging (Polα) strand synthesis offers the opportunity to examine how well coupled DNA synthesis is throughout the genome. As expected, Polε and Polα usage on the leading and lagging stands respectively, for forks moving in the same direction is notably similar. However, reproducible differences in Polε and Polα profiles are detected (e.g. around 26 Mb on chromosome 6 in **Fig. 5a**, experimental replicate: **Supplementary fig. S9**). We interpret this as evidence of uncoupling between leading and lagging strand polymerases^25^. To quantify this, we calculated a separate coupling index (CI) for rightward and leftward moving forks to represent the bias toward either Polε or Polα usage (**Fig. 5b**). If both leading and lagging strand synthesis contribute equally to replication of the duplex DNA, CI = 0. Positive values of CI represent a bias toward leading strand polymerase (Polε) while a negative value represent a bias toward lagging strand polymerase (Polα).

**Fig. 5.**
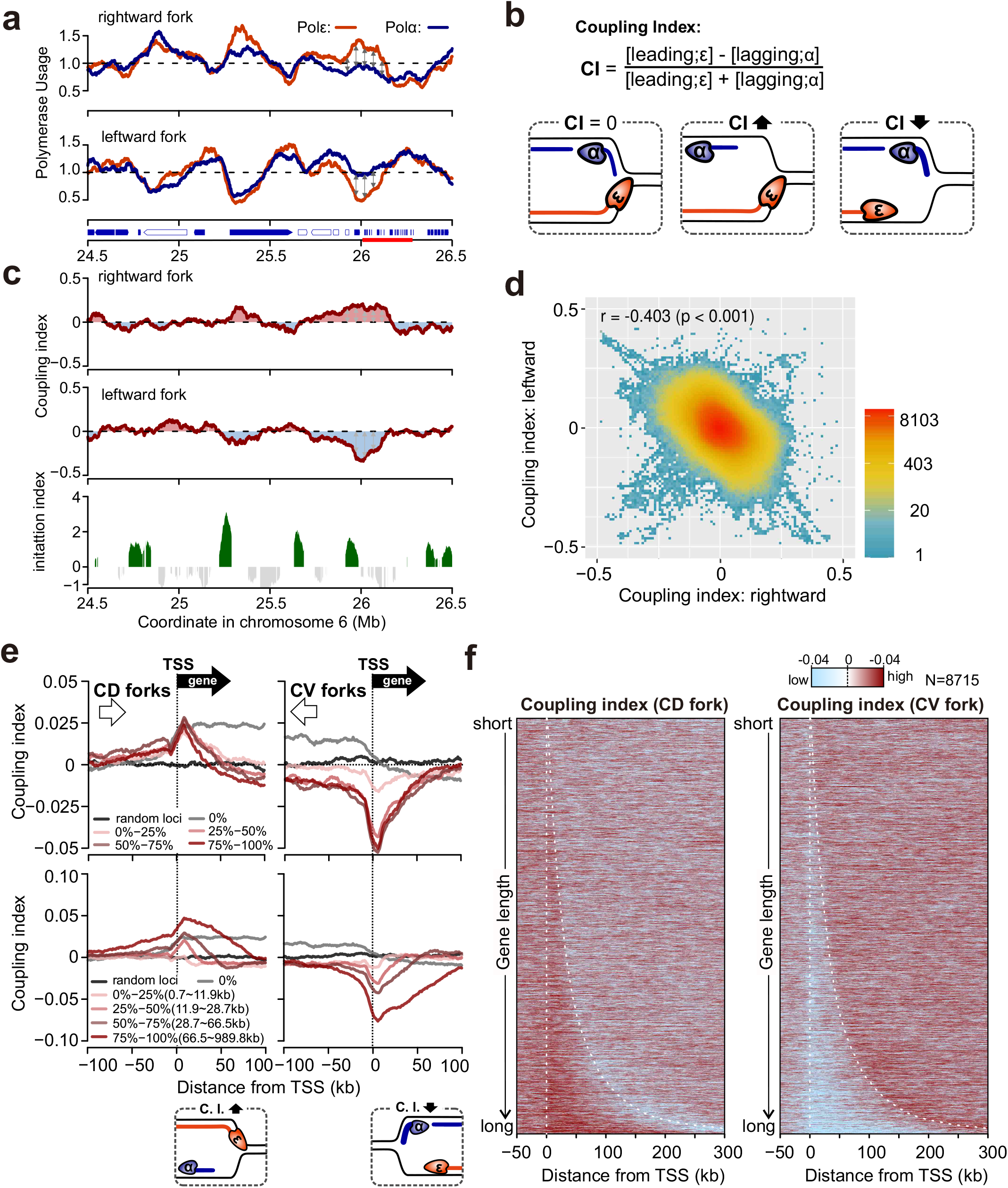
The uncoupling of leading and lagging strand DNA synthesis. **a**. Profiles of polymerase usage for a region of chromosome 6. Arrows on the vertical axis indicate active genes (blue-filled) and inactive genes (unfilled). The red line on the vertical axis indicates a cluster of histone-encoding genes. **b**. Definition of coupling index (CI, for details see materials and methods). **c**. Top: CI of rightward moving forks. Middle: CI of leftward moving forks. Bottom: the initiation index of the same region for comparison. **d**. Correlation of CIs of rightward and leftward forks. **e**. Averaged coupling index +/- 100kb around annotated TSS in the human genome for CV and CD forks categorised by transcriptional activities (top) or gene length (bottom). Only the 50% most transcriptionally active genes were included. **f**. Heat map representation of data in panel e. Data were sorted by gene length. Broken lines indicate the positions of TSS and TTS.

Across the majority of genomic regions the CI remains close to zero, but at some loci it reproducibly deviates by +/-0.25, suggesting up to 25% biased usage of Polε or Polα occurs relatively frequently across the genome (**Fig. 5cd**). We observed a significant correlation between the two biological replicates (**Supplementary fig. 10a**). Interestingly, we observed a reciprocal pattern for CI between rightward and leftward moving forks (**Fig. 5d, Supplementary fig. 10b**). This indicates an opposite bias for forks moving in the two directions is present. For example, at the histone-encoding gene clusters on chromosome 6, usage of Polε was overrepresented compared Polα in the rightward forks and the opposite trend was observed in leftward forks (**Fig. 5c, Supplementary fig. 9**). This inverse correlation is conserved across the genome (replicate 1: r = -0.403 p<0.001, replicate 2: r = -0.354 p<0.001, **Fig. 5d, Supplementary fig. 10b**). We next examined if these CI fluctuations correlated with transcription. Aligning CI of co-directional (CD) and converging forks (CV) at TSS sites we observed that the CI of CD forks increased within gene bodies, whereas the CI of CV forks decreased (**Fig. 5ef**). This likely reflects the orientation of DNA polymerase movement and transcription: synthesis by forks moving in the same direction of RNAPII encounter problems that result in a bias toward Polε (suggesting lagging strand synthesis is impaired), while synthesis by forks moving in the opposite direction to RNAPII result in problems that result in a bias towards Polα (suggesting that leading strand synthesis is impaired). Consequently, DNA synthesis toward opposite to transcription was shown to be impaired in both CD and CV forks (further discussed below).

The genome wide distribution of CI variation tends toward lower values (**Fig. 5d**, **Supplementary fig. 10b**). This suggests that leading strand synthesis is generally more susceptible to spontaneous impediment than lagging strand synthesis. The affected regions are not uniformly dispersed across the genome and tend to favour particular chromosomes (**Fig. 6a**). To identify regions associated with this phenomenon, we statistically identified 27 regions as CI outliner loci, where either rightward or leftward CI values diverged significantly from the population of chromosomal data in both biological replicates (**Table 1**). Among these, 16 loci (59.2%) were associated with large genes. In all but one of these the CI was notably low (CI < -0.3) for forks converging with transcription. One explanation of these results is transcription stress due to head-to-head collisions becomes intense in a subset of large genes and consequently polymerase uncoupling occurs frequently due to an impediment of leading strand DNA synthesis. However, it should not be excluded that an alternative engagement of other polymerases, such as Polδ during replication restart^26, 27^ causes the low CI values.

**Fig. 6.**
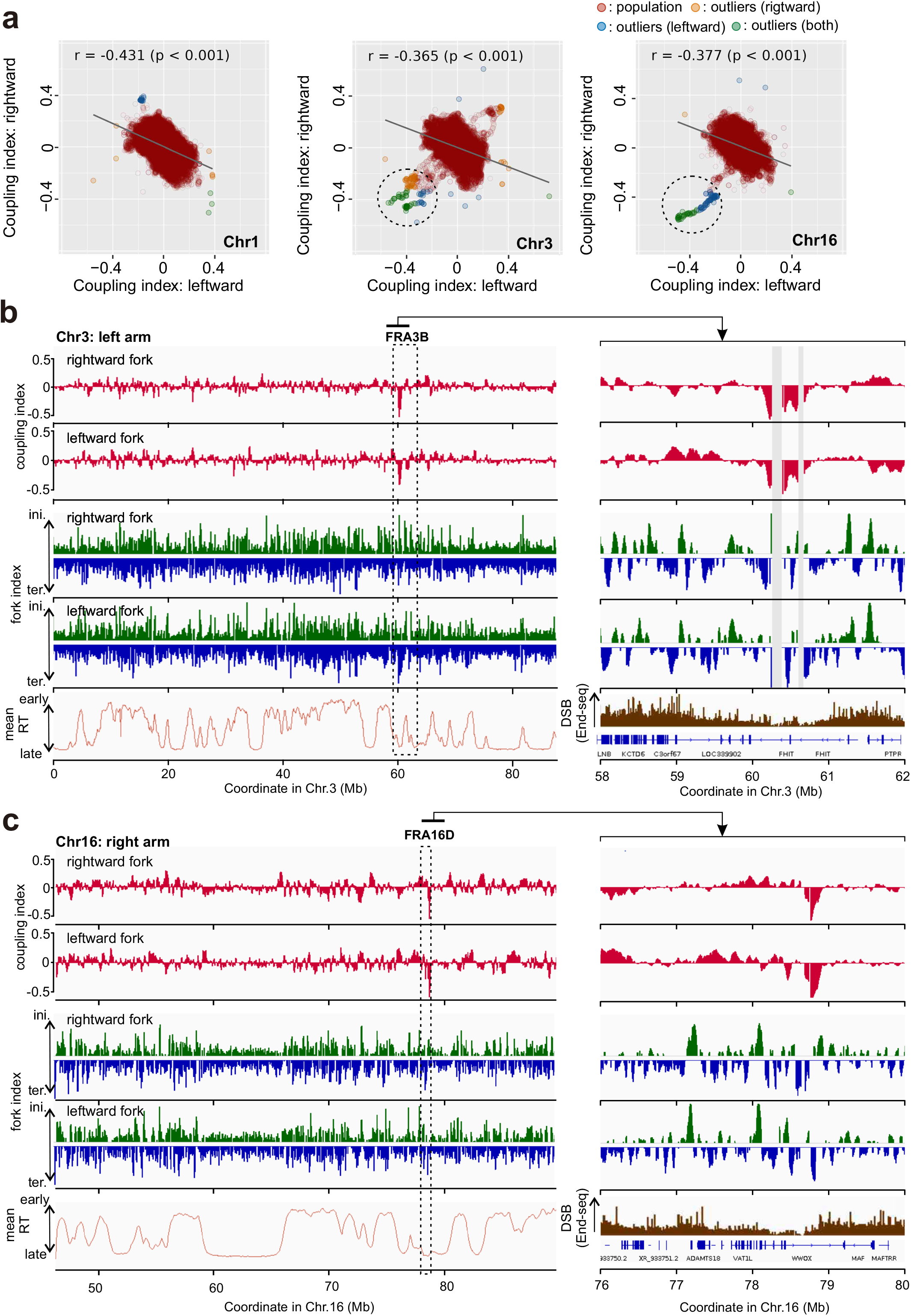
Polymerase uncoupling at common fragile sites FR3B and FRA16D. **a**. Correlation of coupling index of rightward and leftward moving forks presented by chromosome. Chromosomes 1, 3 and 16 are shown. The circles highlight data points dispersed from the bulk of population. These outliner data points were identified by Smirnov-Grubbs test. **b**. Top: profiles of coupling index for rightward and leftward forks. Middle: fork index profile. Bottom: mean replication timing (mean RT) or DSB data for the FRA3B region on chromosome 3. **c**. The equivalent data for the FRA16D region of chromosome 16. Replication timing data for HCT116 cells are derived from Zao et al (2020)^21^. Data of DSBs from End-seq are derived from Tubbs et al (2018)^29^.

**Table 1.** Coupling index (CI) outliner loci

| Chromosome | Position (Mb) | Gene name | Size (kb) | Gene direction | CI <0 ( leading str. < lagging str.)* | CI < 0 ( leading str. < lagging str.)* |
| --- | --- | --- | --- | --- | --- | --- |
| 2 | 235.4-236.0 | AGAP1 | 638 | rightward → | ← leftward fork | - |
| 3 | 60-61 | FHIT | 1520 | ← leftward | ← leftward & rightward → fork | - |
| 3 | 114.4-114.6 | ZBTB20 | 852 | ← leftward | - | rightward fork → |
| 3 | 188.1-189.1 | LPP | 757 | rightward → | ← leftward & rightward → fork | - |
| 4 | 19.5-20.0 | - |  | - | rightward fork → | - |
| 4 | 49.1-49.2 | - |  | - | rightward fork → | - |
| 5 | 36.8-37.1 | NIPBL | 190 | rightward → | ← leftward fork | - |
| 5 | 131.5-131.8 | FNIP1 | 155 | ← leftward | rightward fork → | - |
| 9 | 123.6-124.0 | DENND1A | 550 | ← leftward | rightward fork → | - |
| 10 | 38.4-38.6 | - |  | - | ← leftward fork | - |
| 10 | 41.8-42.1 | - |  | - | ← leftward fork | - |
| 10 | 87.2-87.4 | NUTM2A-AS1 | 139 | ← leftward | rightward fork → | - |
| 13 | 60.0-60.2 | DIAPH3 | 498 | ← leftward | rightward fork → | - |
| 13 | 77.2-77.4 | MYCBP2 | 282 | leftward | rightward fork → | - |
| 13 | 98.1-98.3 | FARP1 | 307 | rightward → | ← leftward fork | - |
| 15 | 41.0-41.2 | INO80 | 137 | ← leftward | rightward fork → | - |
| 16 | 34.6-34.8 | - |  | - | rightward fork → | - |
| 16 | 78.7-78.9 | WWOX | 1113 | rightward → | ← leftward & rightward → fork | - |
| 17 | 21.8-22.0 | - |  | - | leftward fork | - |
| 17 | 26.7-26.9 | - |  | - | leftward fork | - |
| 18 | 49.2-49.6 | DYM | 418 | ← leftward | rightward fork → | - |
| 20 | 31.0-31.3 | - |  | - | rightward fork → | - |
| 21 | 7.9-8.0 | - |  | - | rightward fork → | - |
| 21 | 10.6-10.8 | - |  | - | rightward fork → | - |
| 21 | 19.2-19.6 | - |  | - | ← leftward & rightward → fork | - |
| 22 | 40.4-40.7 | MRTFA | 226 | ← leftward | rightward fork → | - |
| 22 | 47.1-47.2 | TBC1D22A | 433 | rightward → | rightward fork → | - |

\*CI values of the indicated orientation of fork were determined to be outliers in the distribution of the whole chromosome by Smirnov-Grubbs test.

In three of the uncoupling regions (**Table 1**) CI values were significantly negative for both rightward (CD) and leftward (CV) moving forks. This indicates perturbation of leading strand DNA synthesis on both strands of the duplex (**Fig. 6bc**). Two of these three loci are positioned within the FRA3B and FRA16D common fragile sites (CFSs) that are highly expressed in many cell lines, including HCT116^28^. The low CI loci did not correlate with replication initiation or termination zones, suggesting that specific impediments to leading strand polymerases occur at these regions. Comparing our data with published End-seq experiments in HCT116^29^, the low CI loci do not overlap with hotspots of double strand breaks (DSBs). Thus, we propose that uncoupling of replicative polymerase is separate feature of at least a subset of CFS and can contribute to chromosome rearrangement in a manner independent of DSBs.

## Discussion

By locating the positions of incorporated rNMP by mutated DNA polymerases in human HCT116 cells we have characterised genome-wide usage of the replicative polymerases Polε and Polα. The profiles of Polε and Polα usage on either the Watson or Crick strands are strikingly reciprocal. While Polα primes both leading and lagging strand synthesis, the number of lagging strand priming events vasty exceed that of the leading strand. Thus, our data demonstrate that human Polε and Polα contribute to leading and lagging strand DNA synthesis, respectively, across the genome. These results establish that the roles of Polε and Polα, and by implication Polδ, are conserved between human and yeasts^5–7^. Our attempts to characterise Polδ were not successful because the POLD1-L606G or POLD1-L606M mutants (predicted to elevate rNMP-incorporation) could not be isolated.

### Replication initiation

The Polε and Polα profiles for the Watson and Crick strands represent the population average of fork directionality. Our data provide four independent measures of fork directionality. By combining these, we calculated genome wide replication fork directionality (RFD) profiles with high resolution. Sites of frequent replication initiation are, by definition, characterised by changes between the leading and lagging strand polymerases on both the Watson and Crick strands. We therefore calculated an initiation index for each location (1-kb bin), defining locations characteristic of initiation as those where all four polymerase profiles exhibited the expected difference from its neighbouring location. Approximately 12,000 peaks of positive initiation index scores were identified in each biological replicate, a number comparable of those identified by Ori-SSDS for mouse cells (11000 – 13000)^30^. The intra peak distance average was 230 kb, slightly larger than previous estimates (160-190 kb)^31^. The average width of the zones of positive initiation index scores was (33 kb), comparable to OK-seq (mean 30 kb, range 6-150 kb^17^) and optical replication mapping (mean 32 kb^32^). These data confirm that initiation in human cells is zonal.

### Transcription and initiation dynamics

Consistent with previous reports, we observed a correlation between replication initiation and TSS and TTS of transcribed genes. The high resolution of our data allowed us to unambiguously map the centre of these initiation zones ∼20kb upstream and downstream of the TSS and TTS, respectively. While the correlation between TSS/TTS and initiation was highly significant, our data also demonstrate that 20% of genes that are transcribed above the average level do not show increase in initiation enrichment close to TSS/TTS sites and that 60% of initiation zones do not localise in the vicinity of TSS/TTS. By examining chromatin features for correlation with initiation zones, we identified H2AZ and, to a lesser extent H3K27me3, as peaking with initiation. Modifications directly associated with transcription, such as H3K4me3 (TSS associated^23^) and H3K36me3 (gene body associated), showed separate and distinct profiles. These appear to reflect the proximity of the modification at TSS or gene bodies and are likely not causative correlations. We speculate that H2AZ and H3K27me3 contribute to configuring an accessible local environment for initiation zones, in which H2AZ and/or other chromatin features recruit ORC or pre-RC components^33^.

### Uncoupling of leading and lagging strands

A novel aspect of our approach is that having strand specific data for two independent polymerases allowed us to define a fork coupling index (CI – see **Fig. 5b**) that quantifies how well leading and lagging strand synthesis are coupled during fork progression. Surprisingly, a reproducible bias toward either Polε (leading) or Polα (lagging) strand polymerases is relatively common and ranges up to 25% at some loci. The fluctuation of the CI for human cells appear much greater than that previously observed in yeasts^5, 10^ and, unlike in yeasts where polymerase bias is associated with initiation or termination, CI fluctuations are prominent in regions not associated with these phenomena. Molecular events that might generate CI fluctuation include the inhibition of DNA synthesis to generate strand-specific polymerisation defects or DNA synthesis by other polymerases. Thus, CI fluctuation likely reflect multiple mechanisms that can occur either within ongoing replication forks or after fork passage or collapse.

Studies using *E. coli* or budding yeast protein or *Xenopus* extracts^34–36^ showed that, upon uncoupling of the helicase and leading strand polymerase, the helicase slows to encourage recoupling. In vitro analysis using budding yeast proteins suggests that Polδ recouples synthesis. In our analysis, this would manifest as a negative CI (bias towards Polα). Alternatively, if DNA polymerase usage at uncoupled regions is determined by noncanonical polymerases not intrinsic to the fork, the CI would fluctuate toward either positive or negative values, depending on whether leading or lagging stand synthesis was influenced. While noncanonical polymerase usage has mainly been associated with DNA damage tolerance, there is increasing evidence for such phenomena under unperturbed conditions, particularly in mammalian cells^37–41^. Our data thus raise the important question of how extensively the unanticipated level of the plasticity of DNA polymerase usage underpins replication of the human genome. Of note we demonstrate that inhibition of the leading strand polymerase is particularly evident in FRA3B and FRA16D in the absence of exogenous replication stress.

### The influence of transcription on replication fork dynamics

By separately analysing the polymerase usage data associated with rightward (Watson; leading strand template, Crick; lagging strand template) and leftward (Watson; lagging strand template, Crick; leading strand template) moving forks we can evaluate the effect of transcription on the dynamics of either convergent or co-directional replication forks (see **Fig. 4**). This established that, at immediately upstream of TSS, co-directional forks are impaired – most likely when they encounter paused RNAPII, which is frequent around transcription initiation sites. Similarly, convergent forks moving from the gene body showed evidence of impaired processivity immediately downstream of TSS. A similar distinction between converging and co-directional forks was not evident at TTS, where both appear to be impeded upstream of TTS within the gene body. Thus, we reveal distinct fates for codirectional and convergent forks at transcription start sites. To our knowledge, no evidence of perturbation of replication forks at TSS or promoter regions has previously been reported.

The accumulation of RNAPII pausing has been shown to be a phenomenon associated with cancer- prone situations^42, 43^. The consequent increase in replication-transcription conflicts, such as we demonstrate here, may underlie this association. We note that the impairment to fork dynamics calculated for a single direction (Fk^L^ and Fk^R^) does not appear in our initiation indices (Ini^ε^, Ini^α^ and Ini^ε|α^) because initiation indices are defined to describe fork dynamics across both orientations. Thus, by establishing the effects of transcription on both the initiation index and fork index, our analysis separately detects the impediment to fork progression at TSS or the termination of two merging forks, which we show occur at different preferential locations relative to genes^42, 43^.

Consistent with clashes between replication and transcriptional machineries, we observed increased coupling index fluctuations in gene bodies, implying transcription-replication conflicts cause frequent polymerase uncoupling. Separating out the effects on convergent and co-directional forks, the block to DNA synthesis tends to manifest where the non-transcribed strand is used as template DNA. One potential explanation is that, when DNA is transcribed, R-loop structures can form behind RNAPII. These would lead to transient regions of ssDNA on the non-transcribed strand. ssDNA is chemically less stable than dsDNA and more susceptible to damaging agents and some DNA-modifying enzymes^44^. It is also of note that transcription coupled repair eliminates DNA damage specifically on the transcribed strand. As a result, DNA damage on the non-transcribed strand may become relatively more influential in perturbing DNA synthesis. Whatever the underlying causes, this transcriptional effect is particularly manifested in large genes, which may reflect the fact that transcription must proceed beyond the end of S phase^45^. One possibility is that the accumulation of positive supercoiling ahead of, and negative supercoiling behind, RNAPII not only pauses replication forks but also enhances the level of transcription-induced blocks, perhaps by inducing excessive fork rotation, which previous studies in budding yeast have found to generates post-replicative stress^46^.

In summary, Pu-seq in human cells provides a powerful and straightforward methodology to explore DNA polymerase usage and replication fork dynamics. We show here that data produced from Pu-seq are highly consistent with those from other methods such as high resolution repli-seq, OK-seq. The two independent datasets required for Pu-seq provide four separate measures of replication dynamics, allowing predictions of replication fork movement, initiation and termination with great accuracy. Exploiting this we show that transcription influences circa 40% of initiation events and contributes significantly to replication fork uncoupling and subsequent genome instability. Pu-seq will thus provide a useful tool for examining DNA replication by manipulating specific genetic changes in the HCT116 cell lines we have developed, or by constructing the system in different cell lines harbouring distinct developmental or genetic backgrounds.

## Materials and Methods

### Cell culture

All cell lines are derived from HCT116 cells (ATCC, #CCL-247) and are listed in **Table 2**. Cells were cultured in McCoy’s 5A, supplemented with 10% FBS (Gibco, #10437-028), 2 mM L-glutamine, 100 U/ml penicillin and 100 μg/ml streptomycin at 37°C in 5% CO_2_. IAA (Nakarai tesque #19119-61), 5- Ph-IAA (BioAcademia #30-003) and auxinole (BioAcademia #30-001) were dissolved in DMSO to create a 500 mM stock solution (stored at -20°C) and further diluted with media to an appropriate concentration and added directly (1:25 v/v) to the culture medium to achieve the working concentration (IAA 500 μM, 5-Ph-IAA 400 nM, auxinole 100 μM). Doxycycline was dissolved in water to create a 1 mg/ml stock solution (stored at -20°C) and added directly (1:1000 v/v) to the culture media to achieve working solution (1μg/ml).

**Table 2.**
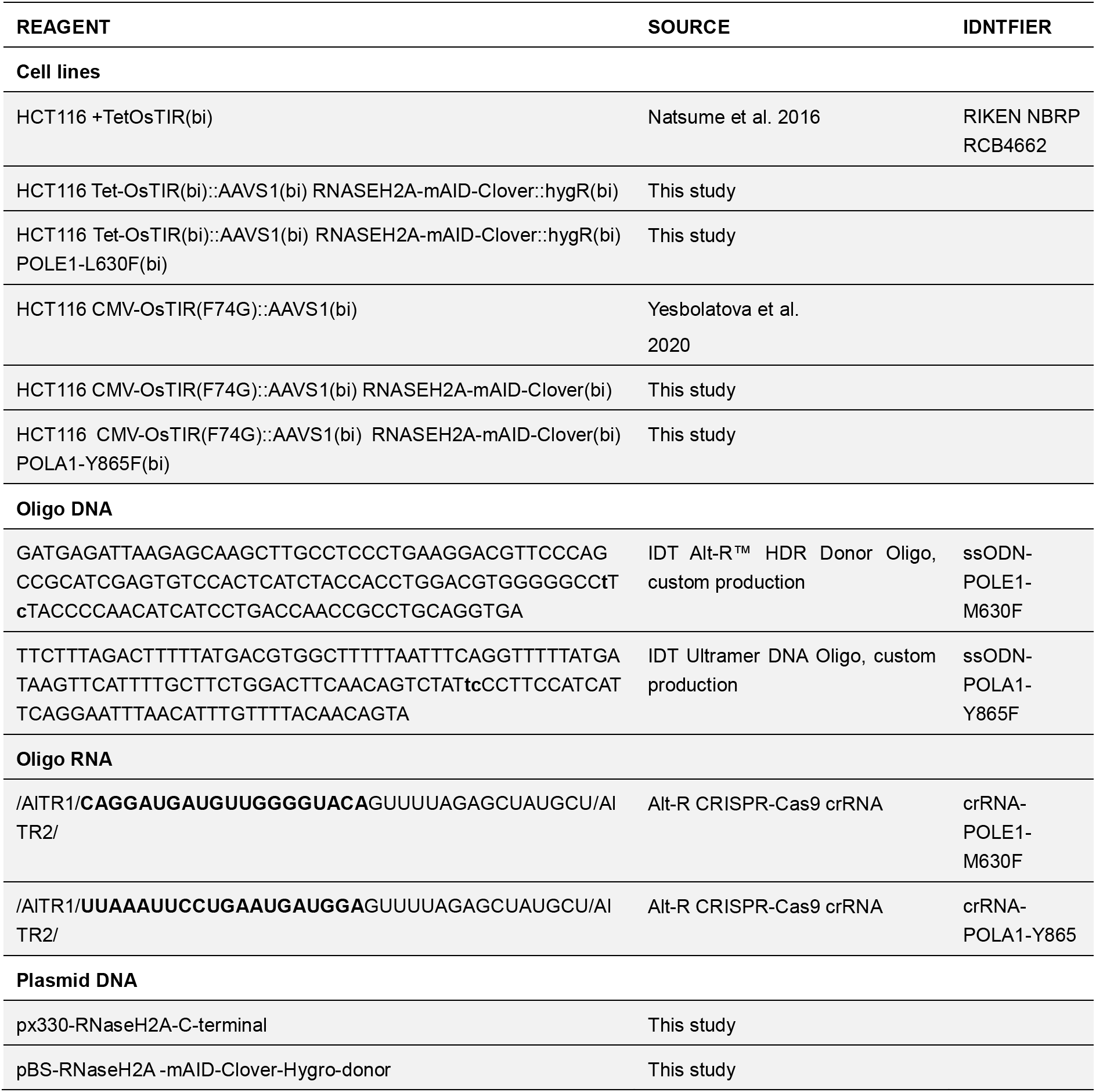
Resource table

### Generation of polymerase mutants

Cas9 protein (Alt-R® S.p. HiFi Cas9 Nuclease V3, #1081060) and tracrRNA (Alt-R® CRISPR-Cas9 tracrRNA, #1072533) and crRNA (Alt-R® CRISPR-Cas9 crRNA, custom production, **Table 2**) were purchased from IDT (Integrated DNA Technologies, USA). Guide RNA (gRNA) was formed by mixing equimolar amounts (50 μM) of tracrRNA and crRNA in duplex buffer (IDT), heating to 96°C for 5 min, and cooling on the benchtop to room temperature. 61 μM of Cas9 and gRNA were mixed at a ratio of 2:3 and incubated for 0.5 -1 hr. Following RNP formation, 1.5 μl of RNP and 1.5 μl of 36 μM single stranded oligodeoxynucleotide (ssODN) template (listed in **Table 2**) was added into 1 x 10^5^ cells in 12 μl of R buffer in Neon® transfection system 10 μL Kit (Invitrogen, #MPK1096) and the cell suspension applied to Neon® transfection system with 10 Neon tips. Following electroporation cells were immediately suspended into 500 μl of media and Alt-R™ HDR Enhancer V2 (IDT, #10007921) to achieve 20 μMwas added. 3 day after transfection, 300-3000 cells were plated in 10 cm dishes for colony formation. 96 single colonies were picked into wells of a 96 well plate. After replicating the clones into another 96 well plate cells were incubated for 2-4 days. The cells on one plate was subjected to genotyping by PCR and those on the other plate was stored at -80°C with Bam banker DIRECT medium (Nippon Genetics, CS-06-001).

### Generation of mAID- Clover–tagged RNASEH2A cell line

Cells were transfected with CRISPR–Cas9 and donor plasmids (**Table 2**) using FuGENE HD Transfection Reagent (Promega, #E2311) in a 6-well plate following the manufacturer’s instructions. Two days after transfection, cells were plated in 10 cm dishes and selected with antibiotics. Selected clones were isolated and confirmed as previously described^47^.

### The degradation of RNASEH2A cell line in AID or AID2 system

For the AID system the +TetOsTIR mAID-Clover–tagged RNASEH2A cells (**Table 2**) were treated with doxycycline and auxinole^47^ for 24 hrs to induce OsTIR1 without background degradation and this medium was replaced with medium containing doxycycline and IAA. The cells were incubated for a further indicated period. For AID2 system, where OsTIR1(F74G) is constitutively expressed, +OsTIR(F74G) mAID-Clover–tagged RNASEH2A cells (**Table 2**) were treated with 5Ph-IAA for the indicated period.

### DNA extraction, alkaline treatment and library preparation

2 x 10^7^ cells were harvested by centrifugation and genomic DNA was prepared using Blood & Cell Culture DNA Midi Kit 100/G Genomic-tips (Qiagen #13343). To examine alkaline degradation 3 μg of DNA was treated with 0.3 M NaOH at 55 °C for 2 hr in 15 μl. The reaction was stopped by adding 3 μl of 1M Tris-HCl (pH7.5). 1 μl of this solution was subjected to TapeStation RNA ScreenTape Analysis (Agilent #5067-5576, #5067-5577, #5067-5578) to detect ssDNA. For library preparation 25 μg of genomic DNA was alkali treated in 0.3 M NaOH at 55 °C for 2 hr, then loaded onto a 1.5% agarose gel and run for 1 hr 40 min at 100 V. The gel was stained with acridine orange (final concentration 5 μg/ml) for 2 hr at room temperature with gentle shaking followed by overnight destaining in water. Fragments of 300–2000 bp were excised from the gel and isolated with a gel-extraction kit (Macherey-Nagel, NucleoSpin Gel and PCR Clean-up, #740609). Library preparation was performed as previously described^5, 48^. Libraries were 150-bp paired-end (PE) sequenced on an Illumina Hiseq X platform (Macrogen, Tokyo, Japan).

### Analysis of Polymerase Usage

For each sample approx. 200 million PE read were obtained. Raw reads were aligned to GRCh38 using Bowtie2 (version 2.3.5). Those which aligned to multiple genomic locations with the same mismatch scores (AS and XS scores as outputted by Bowtie2) were excluded using a custom Perl script: sam-dup-align-exclude-v2.pl (available at the GitHub site: https://github.com/yasukasu/sam-dup-align-exclude). The position of the 5′ end of each R1 read (which corresponds to the 5′ end of ssDNA hydrolysed by alkali treatment) was determined, and the number of reads in 1-kb bins across the genome were counted separately for the Watson and Crick strands using a custom Perl script: pe-sam-to-bincount.pl (available at the GitHub site: https://github.com/yasukasu/sam-to-bincount). This generated the four datasets for the analysis of one polymerase. In case of Polε: at the chromosome coordinate x, N_w_ ^ε^(x), is the count for RNASEH2-mAID POLE1-M630F on the Watson strand; N_c_ ^ε^(x) is the count for RNASEH2-mAID POLE1-M630F on the Crick strand; N_w_ ^+^(x) is the count for RNASEH2-mAID POL^+^ on the Watson strand; N_c_^+^(x) is the count for RNASEH2-mAID POL^+^ on the Crick strand.

The datasets were normalised using the total number of reads: e.g. N’_w_ ^ε^(x) = N_w_^ε^(x)/∑ N_w_^ε^ for the Polε mutant on the Watson strand (where N’ indicates normalisation). These normalised data were used to calculate relative polymerase usage: e.g. E_w_(x) = N’_c_ ^ε^(x)/N’_w_^+^(x) for usage of Polε on the Watson strand; E_c_(x) = N’ ^ε^(x)/N’_c_^+^(x) for usage of Polε on the Crick strand. The equivalent analysis was performed to yield usage of Polα on Watson and Crick strand: A_w_(x) and A_c_(x). When these data were plotted or used for further analysis (below) they were smoothed using moving average of 2m+1, where m is an arbitrary number the value of which is given in the relevant context. Thus, the data point for each bin is an average of 2m+1 bins: the point of origin and the m bins either side.

### Initiation index and fork index

The difference between each neighbouring data point of polymerase usage was calculated as ΔE_w_(x), ΔE_c_(x), ΔA_w_(x) and ΔA_c_(x) with E_w_(x), E_c_(x), A_w_(x) and A_c_(x) which were smoothed using the value m = 30 (genome-wide plot, **Fig. 2c**, **Fig3b** ,**Fig4b**, **Fig. 5c** and **Fig. 6bc**) or m = 7 (the plot of averages or heat map, **Fig. 3c-e** and **Fig. 4d,e**). These differential data were further smoothed by application of a moving average with value m = 15 (genome-wide plot) or m = 7 (the plot of averages or heat map). At each location where all four polymerase profiles exhibit consistent patterns for initiating bidirectional replication forks (ΔE_w_(x)>0 ∩ ΔE_c_(x)<0 ∩ ΔA_w_(x)<0 ∩ ΔA_c_(x)>0) or patterns consistent with the merging of two forks (ΔE_w_(x)<0 ∩ ΔE_c_(x)>0 ∩ ΔA_w_(x)>0 ∩ ΔA_c_(x)<0) an initiation index was defined as: Ini(x) = ΔE_w_(x) - ΔE_c_(x) - ΔA_w_(x) + ΔA_c_(x). This data was subjected to Z-score normalisation (mean = 0, standard deviation = 1) and Z(0) were subtracted to maintain the original + or – information, which represent increased levels of replication initiation and termination in the cell population respectively. Fork index was similarly calculated, but separately for rightward and leftward moving forks (Fk^R^ and Fk^L^). For example, locations of rightward fork initiation are where ΔE_w_(x)>0 ∩ ΔA_c_(x)>0. Similarly, positions of rightward moving fork termination are where ΔE_w_(x)<0 ∩ ΔA_c_(x)<0. At these positions Fk was calculated as Fk^R^(x) = ΔE_w_(x) + ΔA_c_(x) to give the fork index of rightward moving forks. The equivalent was performed for leftward moving forks: location initiation is where ΔE_c_(x)<0 ∩ ΔA_w_(x)<0 and termination where ΔE_c_(x)>0 ∩ ΔA_w_(x)>0). The index for these locations was calculated as Fk^L^(x) = - dE_c_(x) - dA_w_(x). These data were subjected to Z-scores normalisation and Z(0) subtracted.

### RFDs from Pu-seq and Ok-seq

RFD from polymerase usage were calculated by subtraction of polymerase profiles typical of leftward moving fork signals from rightward moving fork signals^17^. When using only Polε usage data, RFD^ε^ = (E_w_(x) - E_c_(x))/ (E_w_(x) + E_c_(x)). When using only Polα usage data, RFD^α^ = (- A_w_(x) + A_c_(x))/(A_w_(x) + A_c_(x)). When using data from both polymerases, RFD^ε|α^ = (E_w_(x) - E_c_(x) - A_w_(x) + A_c_(x))/ (E_w_(x) + E_c_(x) + A_w_(x) + A_c_(x)). To calculate RFD from Ok-seq data, raw sequence read data from OK-seq experiments were obtained from the NCBI Sequence Read Archive (RPE1^18^: SRX4036932, GM06990^17^: SRX1427548, HeLa^17^: SRX1424656) and mapped and counted using the same pipelines as used for the Pu-seq data. Okazaki fragment counts on Watson and Crick strands were defined as OK^W^(x) and OK^C^(x) and the RFD was calculated as: RFD^OK^ = (OK^C^(x) - OK^W^(x))/ (OK^C^(x) + OK^W^(x))^17^. The RFD datasets from Pu-seq and Ok-seq were further smoothed by application of a moving average where m = 3. To convert RFD^ε|α^ to rightward fork proportion, the range between values at -3 standard deviation and +3 standard deviation, which covers 99.7% of data, were converted to 0 to 100%.

### Coupling index

The data of polymerase usage: E_w_(x), E_c_(x), A_w_(x) and A_c_(x) were smoothed using the value m = 30 (genome-wide or 2D plot, Fig. 5c,d, Fig. 6a-c) or m = 7 (the plot of averages or heat map, Fig. 5e,f). To establish a separate coupling index (CI) for both leftward and rightward forks, the lagging strand profile was subtracted from the leading strand profile. For rightward moving forks: CI^R^ = (E_w_(x) - A_c_(x))/(E_w_(x) + A_c_(x)), for leftward moving forks: CI^L^ = (E_c_(x) - A_w_(x))/(E_c_(x) + A_w_(x)).

### RNA-seq data analsys

Extraction of total RNA from HCT116 +TetOsTIR cells was performed by using Monarch® Total RNA Miniprep Kit (New England Biolabs, Inc, #T2010S). cDNA library construction using TruSeq Stranded Total RNA Library Prep Kit with Ribo-Zero Human (Illumina #RS-122-2201) and sequencing on an Illumina NovaSeq6000 platform was performed by Macrogen (Tokyo, Japan). Approx. 21 million pair-end reads were sequenced. Raw sequenced reads were aligned to GRCh38 using STAR (version 2.7.3a, https://github.com/alexdobin/STAR). Annotations of transcript units in the human genome (version 94) were retrieved from the Ensembl Genome Brower (http://www.ensembl.org) and those which are categorised as transcript support level 1 or 2 and Esembl/Havana-merged transcripts (i.e. high probability of correct annotation) were chosen for inclusion in the alignment target set. To obtain data of fragments per Kilobase of exon per Million fragments mapped (FPKM), bam files of mapped reads was analysed using Cufflinks (http://cufflinks.cbcb.umd.edu/).

### Statistics and reproducibility

Two biological replicates were obtained for datasets of Polε and Polα and both were used for all the analysis in this study. Data from one replicate are presented in Figs. 1–6: data from the second replicate showed excellent agreement and where relevant is shown in supplementary data.

### Data availability

All data, including raw sequencing reads and source data for graphs in Figs. 1–6, are publicly available after publication . Custom scripts are available upon request from the corresponding authors.

## Acknowledgements

This work was supported by: JST PRESTO grant (JPMJPR18K7) to YD; JSPS KAKENHI grants (JP16H06151, JP17K19336, JP20H03233 and JP21K19203) to YD; (JP20H05396 and JP21H04719) to MTK; Naito foundation, Takeda Science Foundation, Astellas foundation for research on metabolic disorders and NIG Collaborative Research Program grants (3B2016, 3A2017, 83A2019 and 81A2021) to YD; Wellcome Trust grant (110047/Z/15/Z) to AMC.

## Author contributions

Y.D. conceived the study. Y.D. M.T.K. and A.M.C. designed the experimental approaches. Y.D designed informatics approaches. E.K., Y.K., F.Y., T.N., A.H., T.O. and Y.D. performed experiments and their analysis. Y.D. performed computational analysis. Y.D wrote the manuscript. A.M.C, M.T.K. and T.O. edited the manuscript

## Materials & Correspondence

Should be addressed to Y.D.

---

**Supplementary figure 1.**
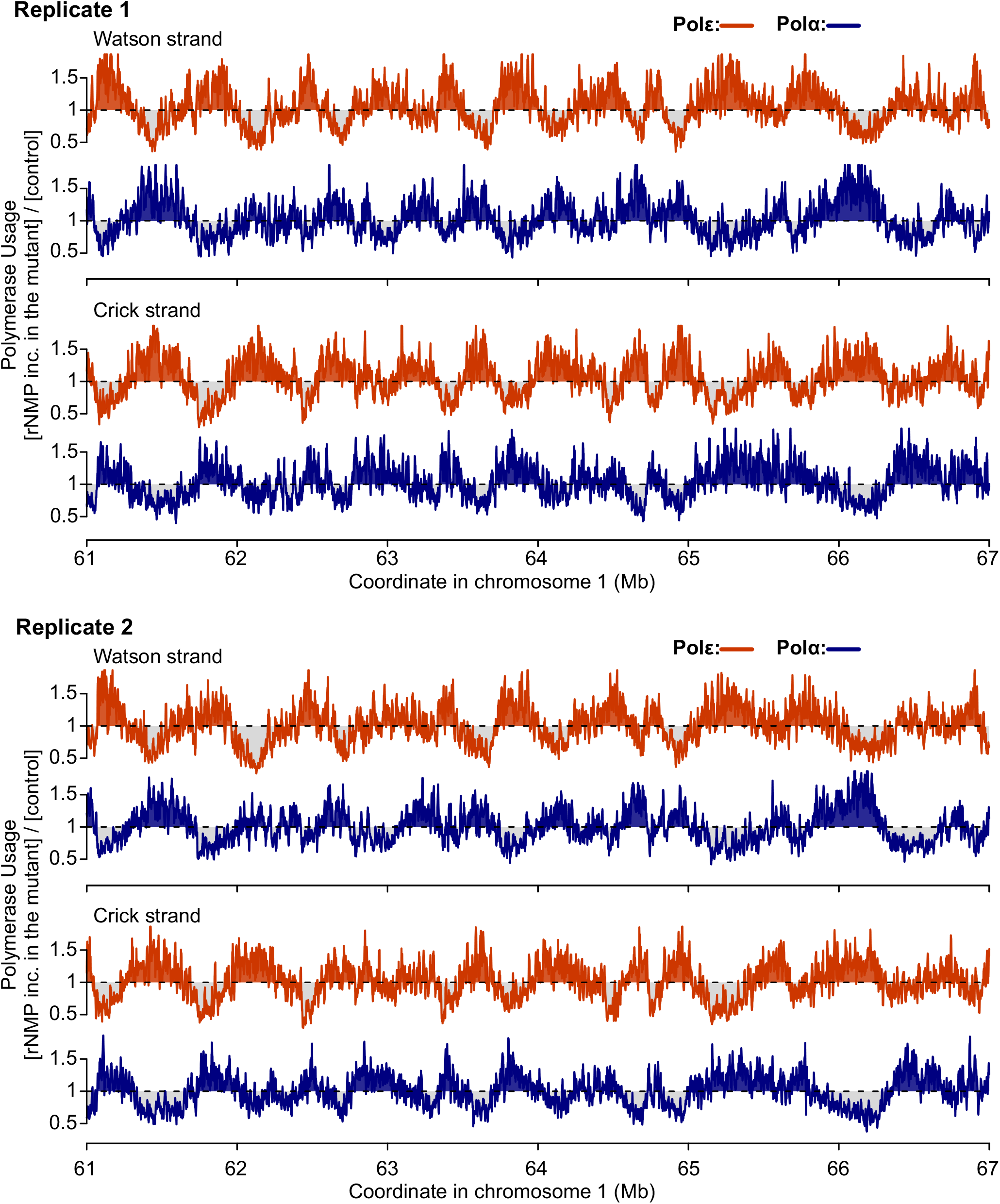
Polymerase usage of 2 experimental replicates across the human genome. For two independent experiments (Replicate 1 and Replicate 2) the profiles of the relative reads for Polε and Polα mutants on the Watson and Crick strand for each 1kb bin are shown. Data are normalised to Pol+. Orange: Polε (POLE1-M630F). Blue: Polα (POLA1-Y865F). A representative region of chromosome 1, a different location to that in Fig. 2, is shown. Data were smoothed with a moving average (m = 3; see materials and methods).

**Supplementary figure 2.**
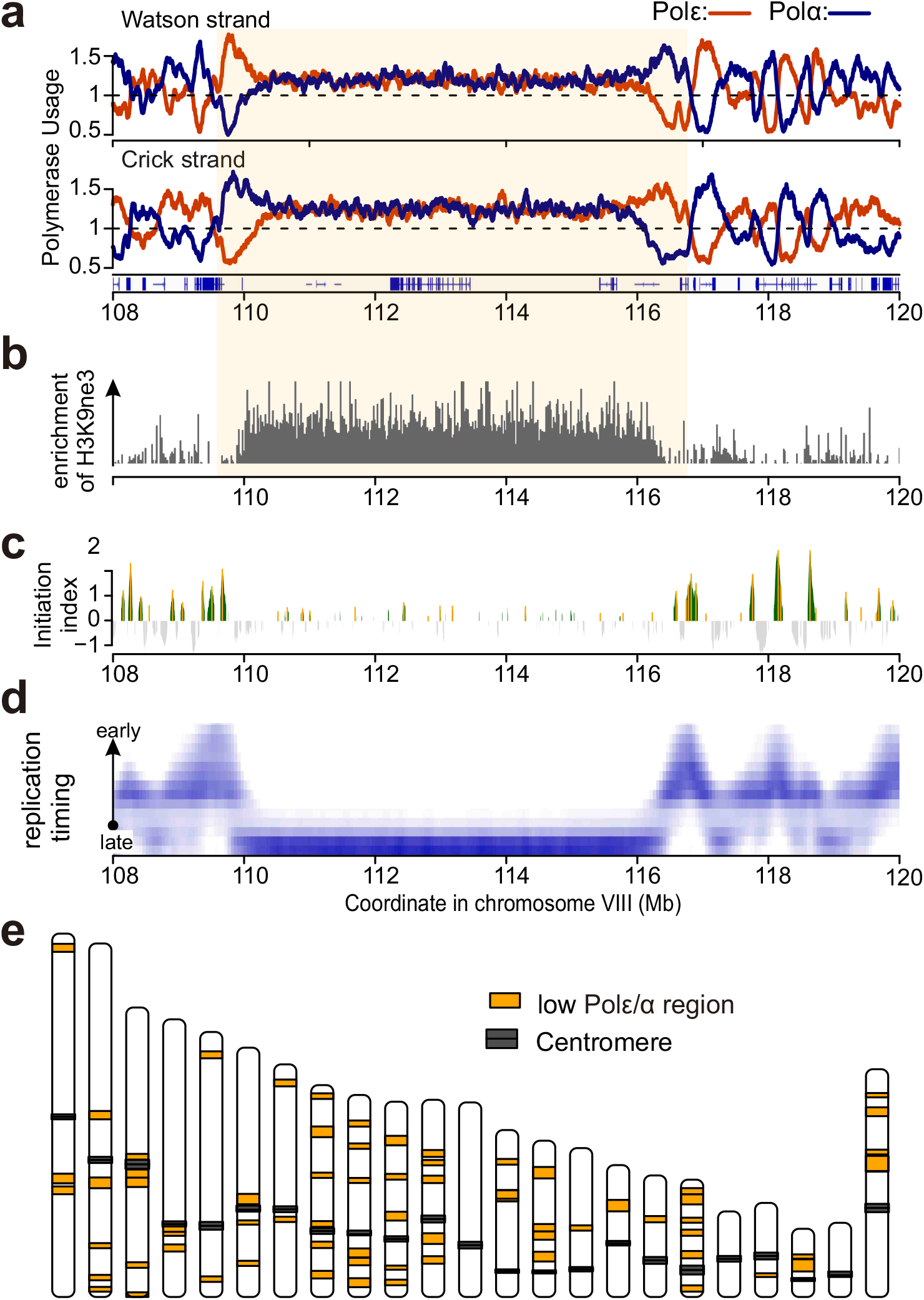
Polymerase usage and replication initiation around late replicating regions. **a**. Polymerase usage profiles of Polε (orange) and Polα (blue) plotted for a representative region showing an equal profile of polymerase usage. Light orange background represents such a ’low Polε*/α* region’. **b**. H3K9me3 enrichment at the same region. Data are taken from Lay et al (2015)^49^. H3K9me3 marks heterochromatin. **c**. Initiation index plotted for the same region. **d**. Replication timing in HCT116 cells. RT data are taken from Zao et al (2020)^21^. **e**. Schematic of the genome-wide distribution of ’low Polε*/α* regions’.

**Supplementary figure 3.**
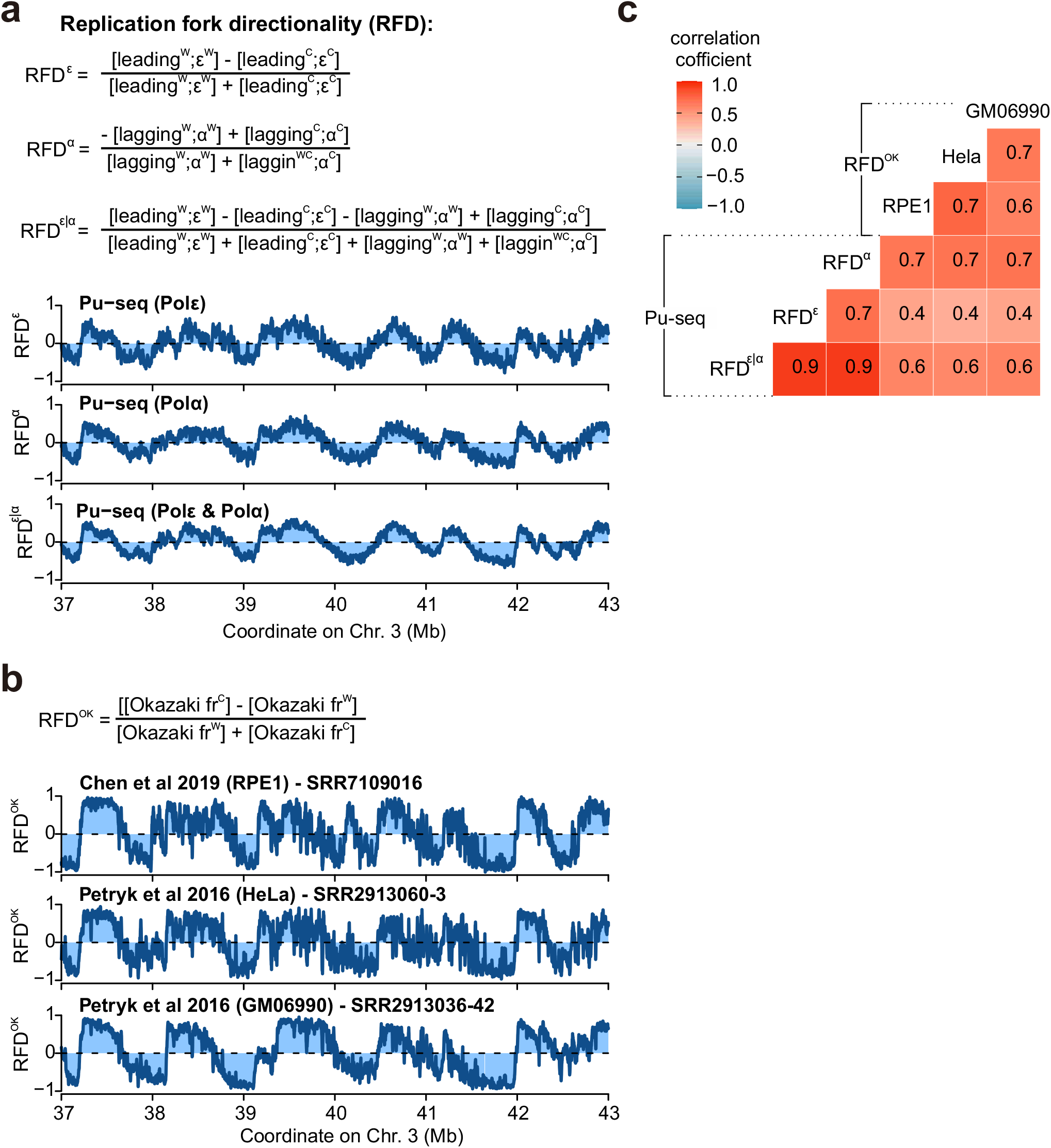
Replication fork directionality (RFD) from Pu-seq and OK-seq. **a**. RFDs are calculated from Polε (RFD*^ε^*) Polα (RFD*^α^*) or both data sets (RFD*^ε^*^|*α*^) and plotted for a representative section of chromosome 3. **b**. RFDs are calculated from three published OK-seq datasets and plotted for the same representative region. **c**. Correlation among RFDs from Pu-seq and OK-seq.

**Supplementary figure 4.**
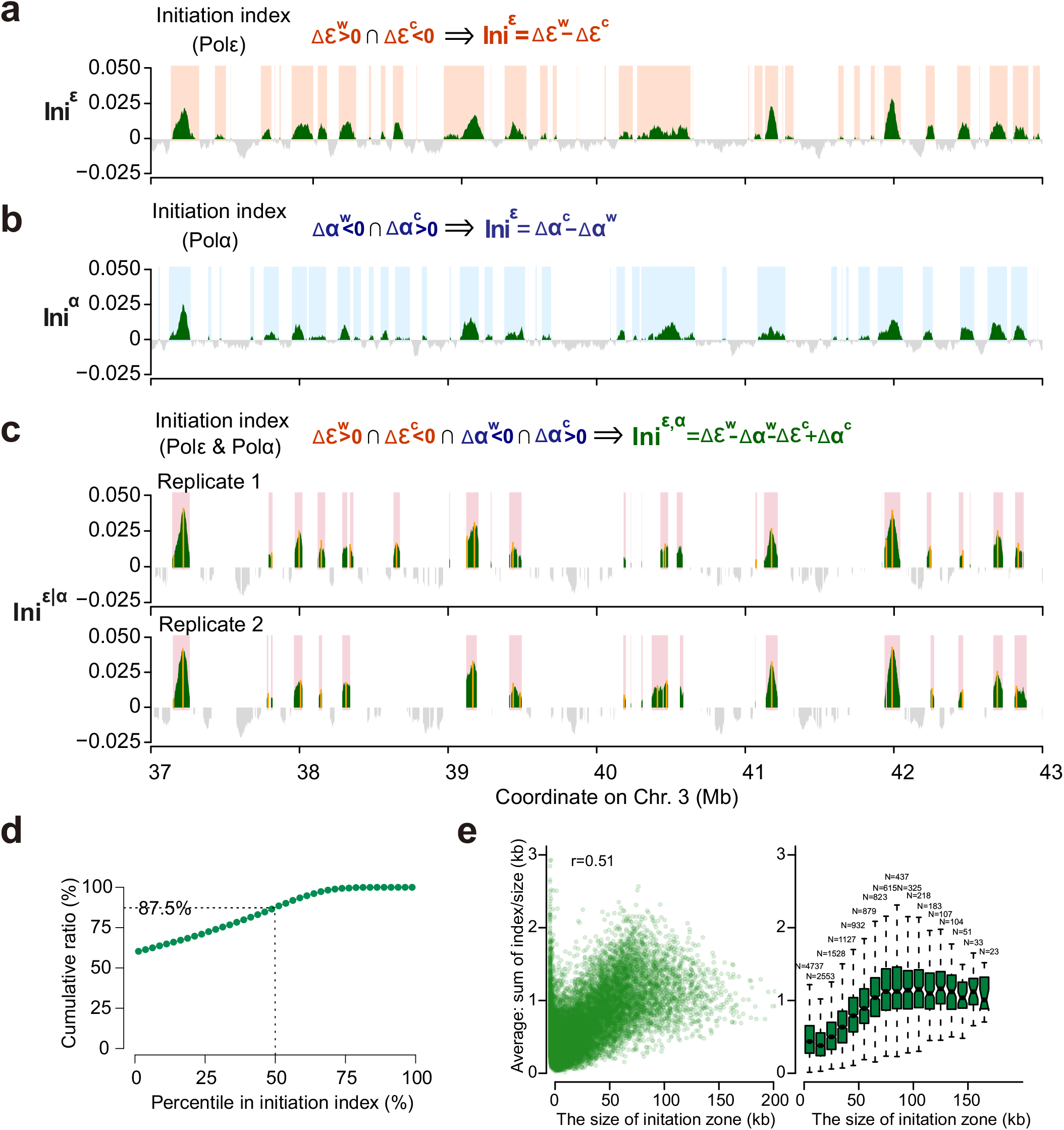
Genome-wide initiation index identifies replication initiation zones. **a.** Initiation index calculated from Polε and plotted for a representative region of chromosome 3. Replication initiation sites are defined as the positive zones of initiation index and are marked with a coloured background. **b**. Initiation index calculated from Polα data and plotted as in panel a. **c**. Initiation index calculated from combined Polε and Polα data and plotted as in panel a. The two independent experimental replicates are shown. Yellow vertical lines indicate computationally identified location of peaks within the zone. **d**. Cumulative ratio of replication initiation sites which locate at the concordant regions between the two experimental replicates. **e**. The relationship of initiation index with the width of initiation zones.

**Supplementary figure 5.**
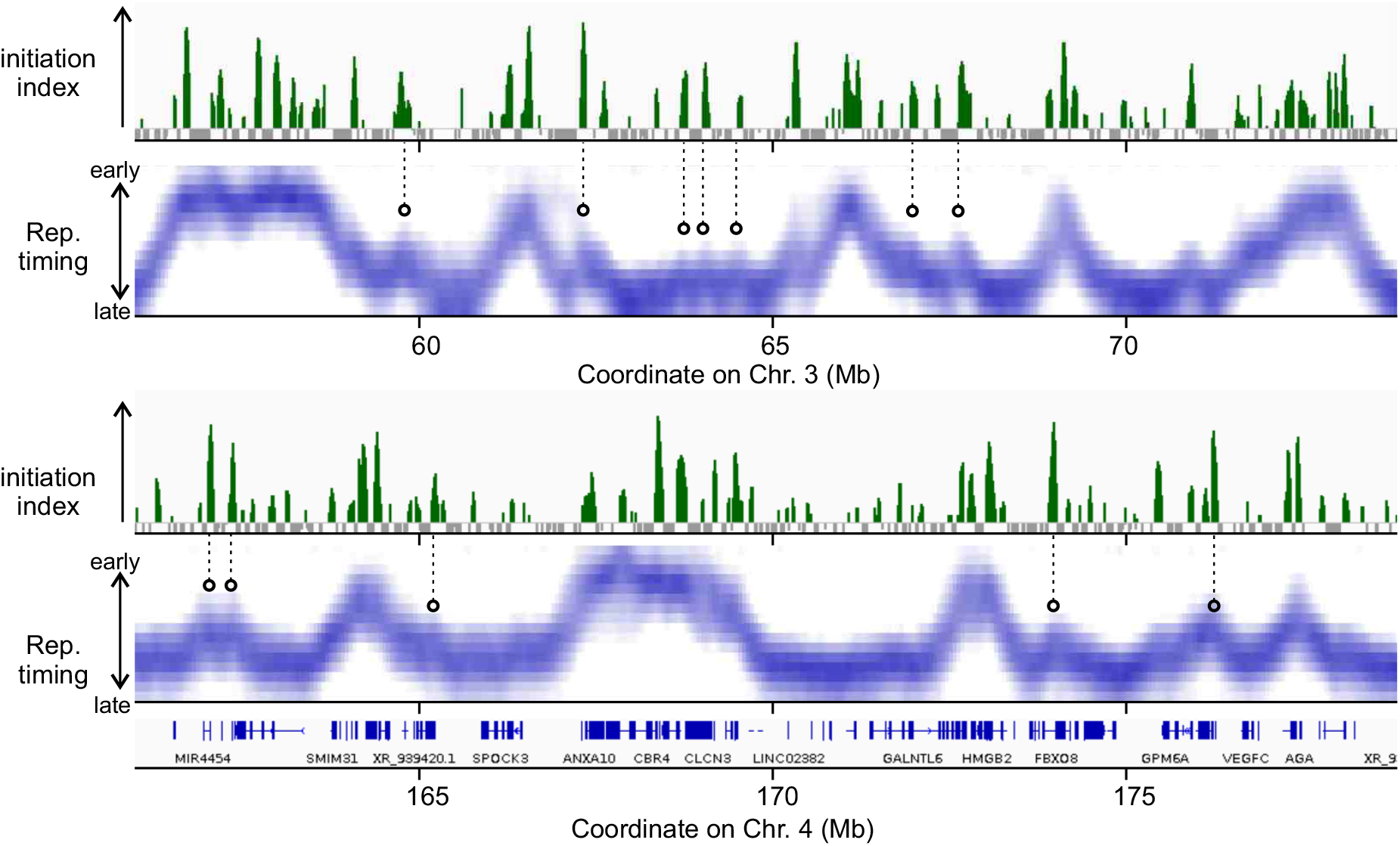
Genome-wide initiation index compared to replication timing. High-resolution replication timing data and initiation index plotted for a representative region of chromosome 3 (top) and chromosome 4 (bottom). The profile of high resolution replication timing for HCT116 cells are from Zao et al (2020)^21^.

**Supplementary figure 6.**
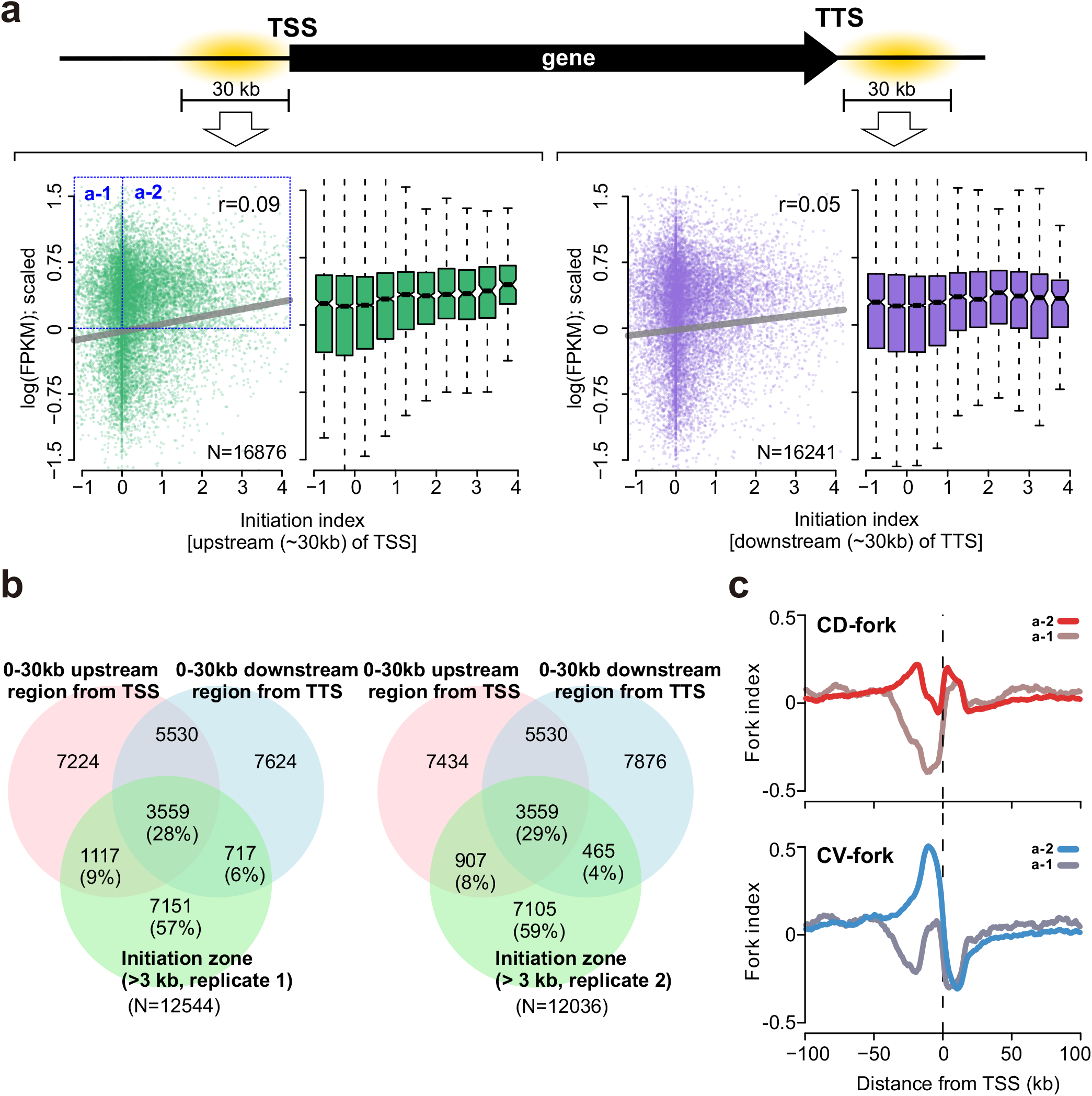
Correlation of initiation and transcriptional activity. **a**. Top: Schematic of a gene showing TSS and TTS. Bottom: Sum of initiation index within the 0-30 kb region upstream of TSS (left two panels) and downstream of the TTS (two right panels) are plotted against transcriptional activity for each gene, represented as Fragments Per Kilobase of exon per Million mapped reads (FPKM) from an RNA-seq experiment (this study). Sum of initiation index and log(FPKM) are normalised to Z-score (average = 0, standard deviation = 1). **b**. Venn diagrams representing the overlap between initiation zones, 0-30 kb upstream of TSS and 0-30 kb downstream of the TTS. **c**. Fork index across a +/- 100 kb around annotated TTS in the human genome. Data of for the fork index around genes with > average transcriptional activity are categorised into two groups as shown in (a): a-1, in which the sum of initiation index in the upstream of TSS is less than or equal to 0: average value (n = 2804), and the a-2, in which this value is greater than 0 (n = 11155).

**Supplementary figure 7.**
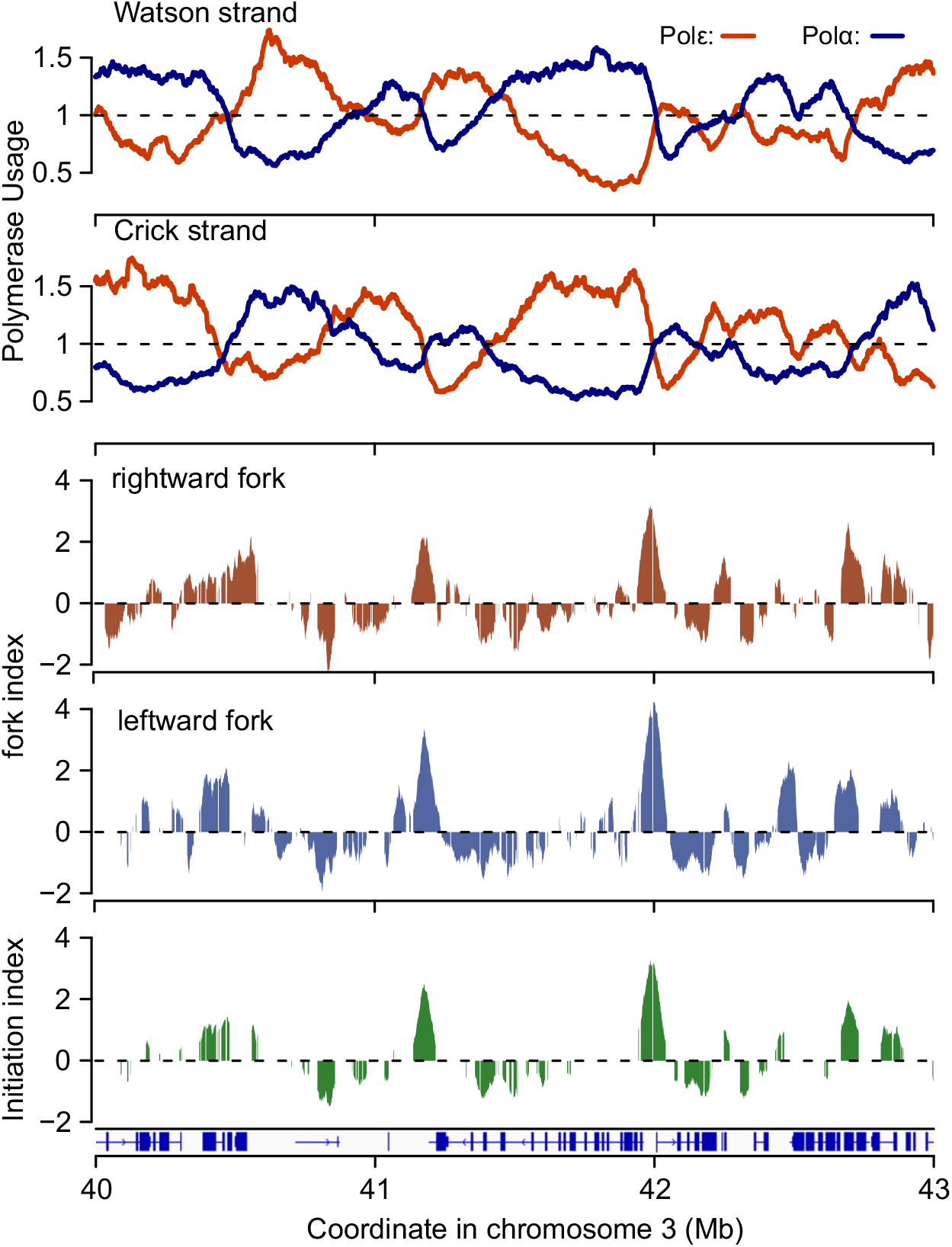
The profiles of rightward and leftward forks in replicate 2. Top: Profiles of polymerase usage plotted for a representative region of chromosome 3. Middle: leftward and rightward moving fork index (Fk^R^ and Fk^L^) plotted for the same region. Bottom: initiation index plotted for the same region. Data used in these plots are derived from the experimental replicate 2 and are comparable to those from experimental replicate 1 that are used in Fig. 4b.

**Supplementary figure 8.**
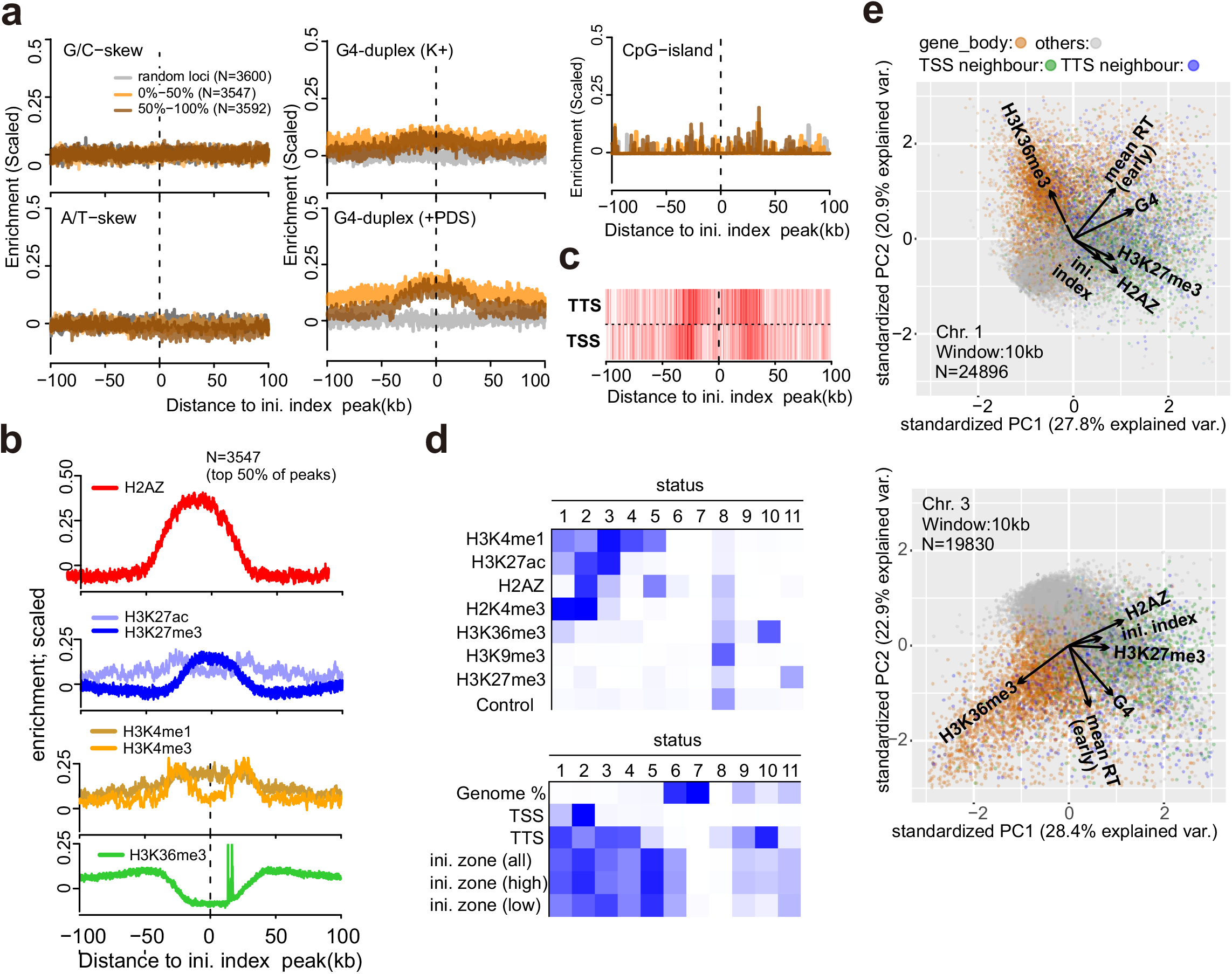
Sequence features and chromatin status at replication initiation zones. **a**. Enrichment of sequence-based features G/C-skew, A/T-skew, G4-duplex and CpG-islands +/- 100 kb around the computationally detected peaks (yellow vertical lines in Extended Data Fig. 4d) in initiation zones that are concordantly detected in the two experimental replicates. Data for genome-wide G4-duplex formation are derived from G4-seq published in Marsico et al (2019)^50^. K+; G4 duplexes formed in the presence of potassium ions. +PDS; G4 duplex formed in the presence of G4-targeting small molecule pyridostatin (PDS). **b**. Equivalent analysis for H2AZ, H3K27 acetylation (ac), H3K27 tri-methylation (me3), H3K4 mono-methylation (me1), H3K4me3 and H3K36me3. Histone modification data are derived from ChIP-seq data for HCT116 published in Lay et al (2019)^49^ and these genome-wide data were z-score normalised before averaging around peaks of initiation index. **c**. Distribution of TSS and TTS around peaks of initiation index. **d**. Heat maps for the model of chromatin-state annotation produced by ChromHMM as described in Ernst and Kellis (2012)^24^. Top: the model was trained on the datasets indicated by the chromatin modification and defined 11 genomic status bins (status 1-11) that represent different chromatin states. The relative enrichments of chromatin features that are present in each status bin are represented by the extent of blue shading (for example, chromosome status bin 10 is enriched only for H3K36me3, which distinguishes it from the remaining 10 bins). Bottom: the model then allocates the locations associated with the TSS, TTS and initiation zones to the same 11 bins (status 1-11), again representing the relative % occupancy by shade of blue (for example, most TSS are associated with bin 2. Referring back to the top heatmap reveals the correlating chromatin marks). **e**. Principal component analysis to generate a correlation map of initiation index scores with the indicated genomic features. Top: chromosome 1. Bottom: chromosome 3. The arrows approximate the direction of each feature and their lengths approximate variances of the data.

**Supplementary figure 9.**
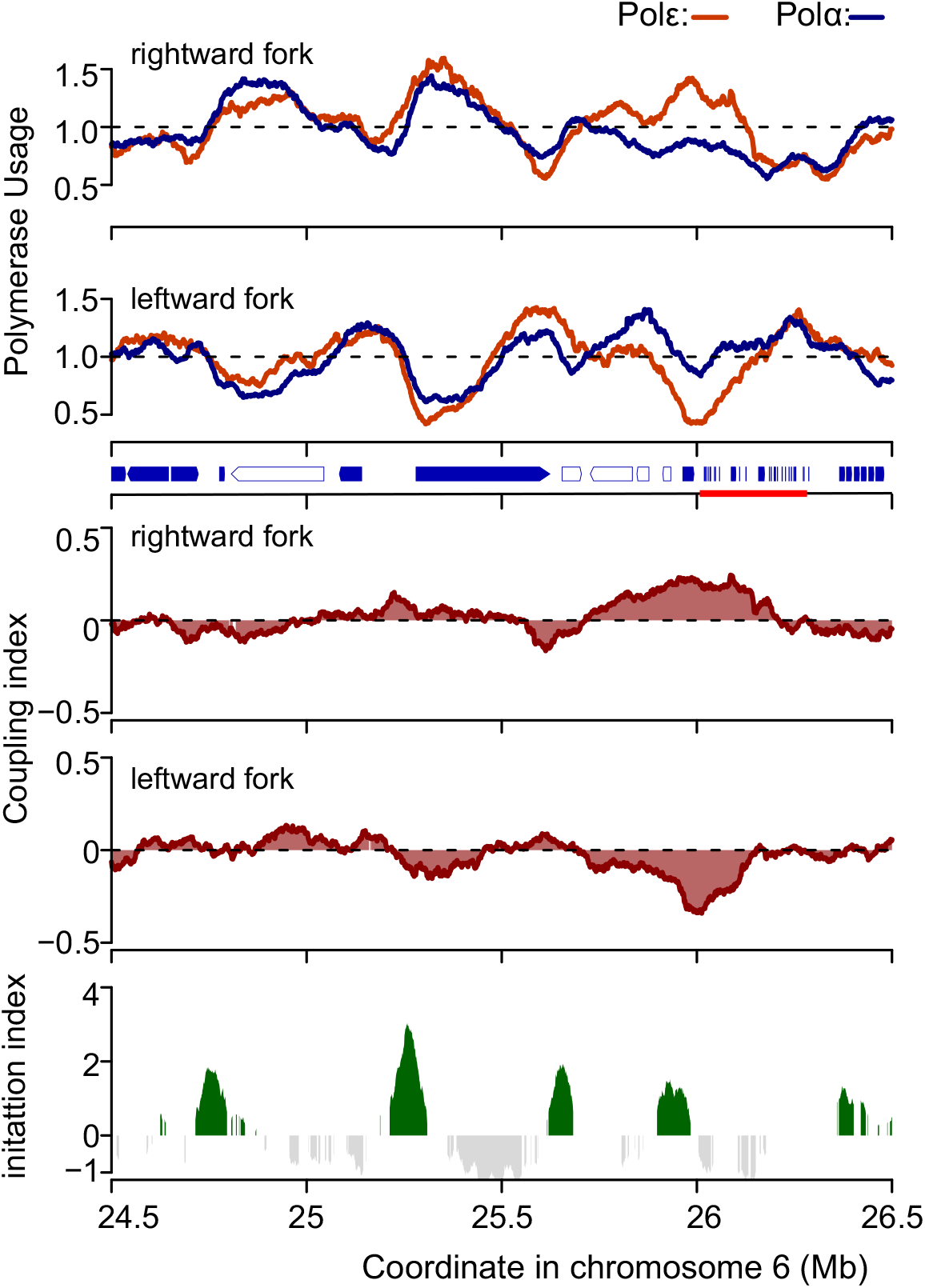
DNA polymerase coupling/uncoupling in replicate 2. Equivalent data for replicate 2 as presented across the same representative region of chromosome 6 for replicate 1 in Fig. 5a,c. Top: Profiles of polymerase usage. Arrows on the vertical axis indicate active genes (blue-filled) and inactive genes (unfilled). The red line on the vertical axis indicates a cluster of histone-encoding genes. Middle: coupling index (CI) of rightward and leftward moving forks. Bottom: the initiation index of the same region for comparison.

**Supplementary figure 10.**
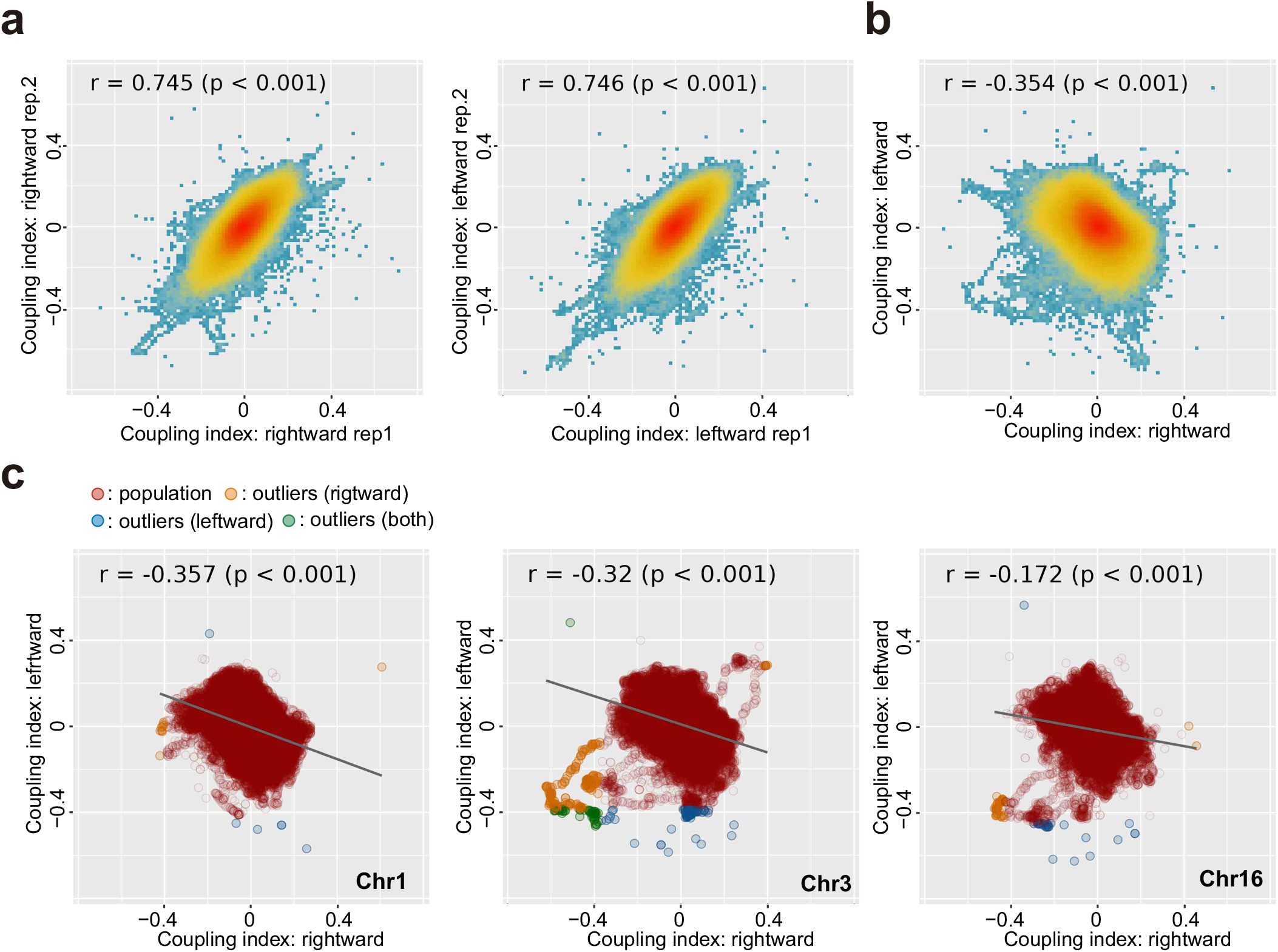
The distribution of coupling index. **a**, correlation of coupling index (CI) derived from two experimental replicates. left: rightward moving forks, right: leftward moving forks. **b**. Correlation of CI of rightward and leftward moving forks from replicate 2. See Fig. 5d for equivalent analysis for replicate 1. **c**. Correlation of CI of rightward and leftward moving forks from replicate 2 presented by chromosome. Chromosomes 1, 3 and 16 are shown. Equivalent data for replicate 1 are shown in Fig. 6a.

